# Dynamics of the Previte-Hoffman food web model with small immigrations

**DOI:** 10.1101/332387

**Authors:** Sk. Sarif Hassan

## Abstract

The ecological dynamics of prey-predator systems are extensively studied over a century. *Previte and Hoffman* introduced a three species(predator-prey with an omnivore) model in 2013. We demonstrate the asymptotic stability of the Previte and Hoffman modified model adding with small immigration into the prey or predator or omnivore population. A comparison of dynamics is also drawn with immigration and without immigration. It is worth noting that the qualitative dynamics is unchanged among the immigration system over the classical Previte-Hoffman system.

## 1 Introduction

The theoretical studies of the prey predator model(well known Lotka-Volterra model) dynamics began by Alfred Lotka and Vito Volterra independently developed in the 1920s [1]. In recent days, there are plenty of works related to qualitative and quantitative dynamics of population/ecological models aim on the study of three-species prey-predator systems [2, 3, 4]. This is possibly because of the chaotic dynamics exhibited by the three-species models as reported by Hastings and Powell [5, 6, 7, 8, 9, 10, 11]. In those models where chaos has been seen, the functional responses of the intermediate consumer and the top predator were assumed to be nonlinear and saturating [12]. A very close look in nature to the one presented here can be found in [12] where a nice introduction to such omnivory model can be relished. Tanabe and Namba [12] numerically demonstrate that the addition of an omnivore(defined as feeding on more than one trophic level leads to a Hopf bifurcation and period doubling cascades. A quick literature review can be obtained from the articles which are studied by various authors [13, 14, 15, 16, 17, 18, 19, 20, 21, 22, 23, 24, 25, 26, 27].

An omnivore is defined as a predator feeding on more than one trophic level and it was commonly observed in some three or more-species food chain models. The omnivore considered in this work is introduced as a scavenger top-predator, which does not only consume the carcasses of the predator but also predates the original prey [28]. Previte and Hoffman introduced a model that describes the dynamics of the predator-prey model with an omnivore [29, 30]. It is shown that this model has the biologically desirable property that all orbits remain bounded. The model exhibits sometime chaos and has infinitely many bounded paired cascades and at most finitely many unbounded cascades.

The paper is organized as follows. In the next section, different possible immigration over three different populations are introduced. Then local stability analysis(mostly numerically) have been shown for each case. Next section sums up a comparison among all the immigration models over the classical Previte-Hoffman model. At last a concussion has been made with possible future endeavours.

In this article, we have started with the original food web model which comprises prey(*x*), predator and a third species(*z*), which consumes the carcasses of the predator along with predation of the original prey. We then modify this model with addition of small immigration into the prey, predator and omnivore populations. We intend to understand the dynamics of the model with immigration and without immigration into the populations.

The original system by *Previte and Hoffman* is described as [29, 30]:

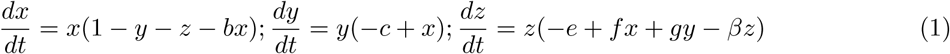

This is a three-species predator-prey model, where the third species is omnivore. The *x, y* and *z* are, respectively, the density of predator, prey, and omnivore populations as functions of time. All the parameters *b, c, e, f, g* and *β* are assumed to be positive. The parameters are interpreted as follows: *b* is the carrying capacity of the prey *x*, *c* is the death rate of the predator *y* in the absence of prey *x*, *e* is the death rate of the omnivore *z* in the absence of its food *x* and *y*, *f* is the efficiency that *z* preys upon *x*, *g* is the degree of efficiency that the omnivore benefits from carcasses of predator *y* and finally *β* is the carrying capacity of *z*.

we have considered a predator-prey systems by adding an immigration factor *p*(*x*) into the prey population or adding an immigration factor *q*(*y*) into the predator population and adding an immigration factor *r*(*z*) into the omnivore population in the *Previte and Hoffman* model as described in Eq.(1).

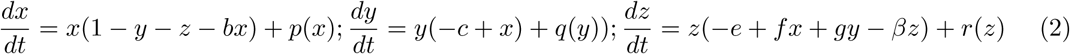

Here *p*(*x*) = *p*, *q*(*y*) = *q* and *r*(*z*) = *z* are considered to be a small real numbers in the interval(0, 1) and *p, q* and *r* represent the number of prey, predator and omnivore immigrants.

Here, we consider the following different possible cases of immigration factor to investigate its effect to the long-term population dynamics of the system Eq.(2).

- Case *I*11: prey immigrants(i.e., *p*(*x*) = *p, q*(*y*) = 0, *r*(*z*) = 0)
- Case *I*12: predator immigrants(i.e., *p*(*x*) = 0, *q*(*y*) = *q, r*(*z*) = 0)
- Case *I*13: omnivore immigrants(i.e., *p*(*x*) = 0, *q*(*y*) = 0, *r*(*z*) = *r*)
- Case *J* 11: prey immigrants relative to the density of prey(i.e., *p*(*x*) = *px, q*(*y*) = 0, *r*(*z*) = 0)
- Case *J* 12: predator immigrants relative to the density of predator(i.e., *p*(*x*) = 0, *q*(*y*) = *qy, r*(*z*) = 0)
- Case *J* 13: omnivore immigrants relative to the density of omnivore(i.e., *p*(*x*) = 0, *q*(*y*) = 0, *r*(*z*) = *rz*)

It is worth mentioning that without loss of any generality we assume the *p, q* and *r* as 0.01 i.e. 1% immigration into the populations are considered throughout this study. Therefore the iterative system of model described in the Eq.(2):

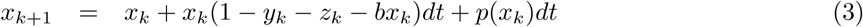

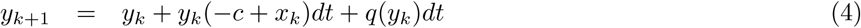

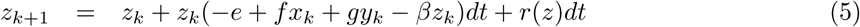

## 2 Stability of the Fixed Points of the Model

In this section we shall describe the local stability of the fixed points of the model for all the cases as stated in the previous section. It is noted that we are only looking for only nonnegative fixed points of the system in order to preserve the biological relevance. The fixed points can be obtained by setting the derivatives Eq.(2) equal to zero [31, 32]. In order to understand the local behaviour of the fixed points one needs to determine the signs of the real parts of the eigenvalues of the Jacobian evaluated at the fixed point. The following theorem is useful for checking the signs of the real parts of the eigenvalues of a 3 × 3 matrix.

### Theorem 2.1. Routh-Hurwitz Criterion

*The Routh-Hurwitz test applied to a general third degree polynomial*

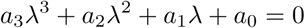

*states that the number of sign changes in the sequence* {*a*_3_, *a*_2_, *H, a*_0_ }*where H* = *a*_2_*a*_1_ − *a*_3_*a*_0_ *is equal to the number of roots of the polynomial having positive real part, and if all entries in the sequence are nonzero and of the same sign, then all roots have negative real part [33].*

### 2.1 Stability of the Fixed Points of the System in the Case *I*11

The system Eq.(2) has four non-negative fixed points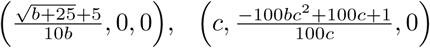,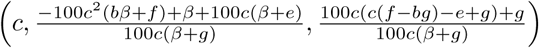 and 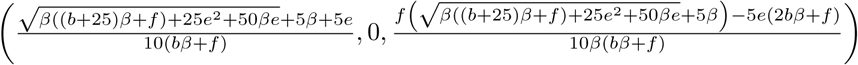.

Here the Jacobian evaluated at the above fixed point say(*x*^*\**^, *y*^*\**^, *z*^*\**^) has the form

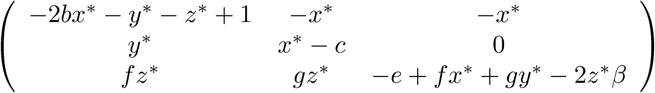

The Jacobian about the fixed point 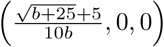is

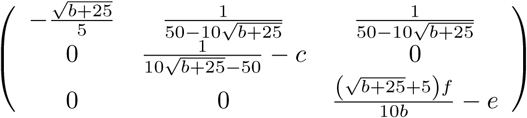

and it has three eigenvalues 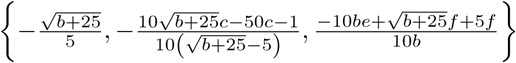.

#### Theorem 2.2.

*The fixed point* 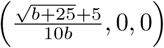 *is locally asymptotically stable if*

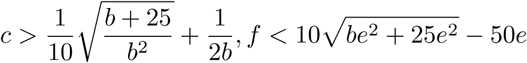

*Proof.* It can be easily seen by doing a simple algebraic calculation that the eigenvalues are negative provided

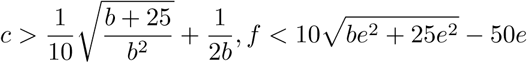

satisfied. Then the fixed point is locally asymptotically stable.

Here we illustrate some example of stability and unstablity of the fixed point 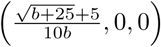. For *b* = 0.616, and for all other positive parameters, the fixed point becomes(1.6333, 0, 0) which is locally asymptotically as shown in Fig.1(left). In addition, due to small non-zero perturbation in the initial predator and omnivore population the fixed point(1.6333, 0, 0) is becoming unstable as shown in Fig.1(right).

**Figure 1:**
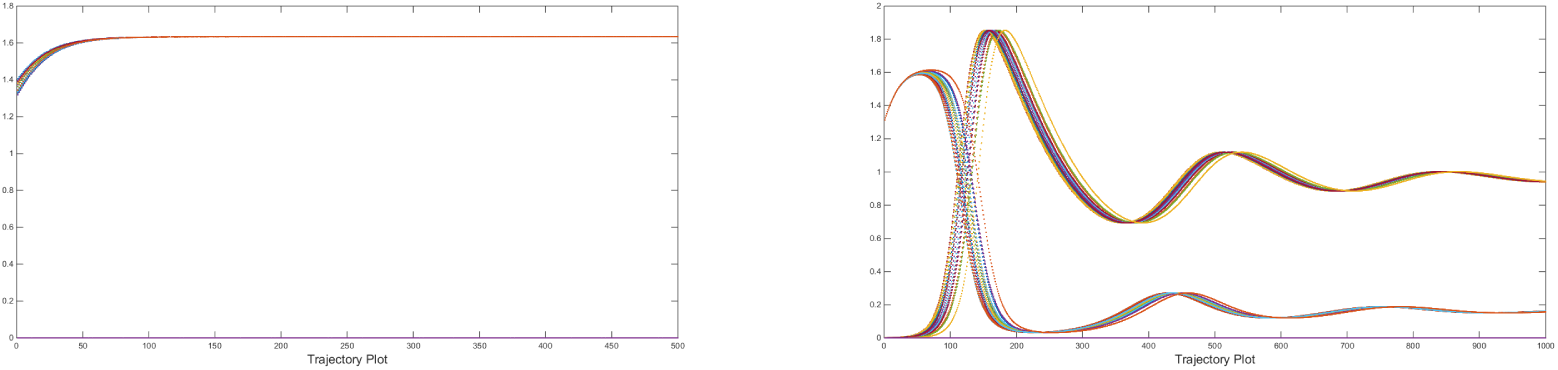
Left: Locally asymptotically stable trajectory of the fixed point(1.6333,0,0) where as in Right: unstablity is seen of the same fixed point due to non-zero initial value of the *y* and *z*. In both the cases 20 different initial values(20 different set of colors are presented in the figure) are taken about the neighbourhood of the fixed point.

The Jacobian evaluated at the fixed point 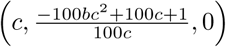 is

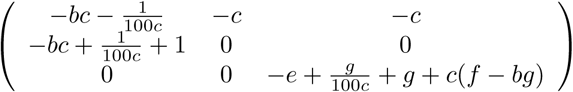

where *a*_3_, *a*_2_, *a*_1_ and *a*_0_(coefficients of the characteristic polynomial) are given as: 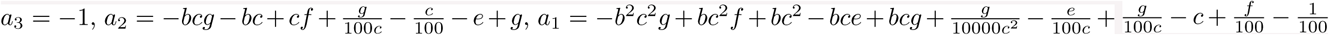and 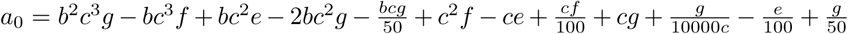 where 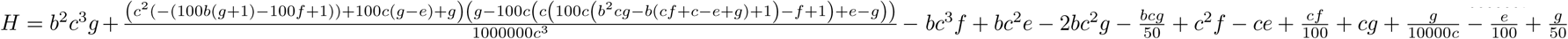

#### Theorem 2.3.

*The fixed point* 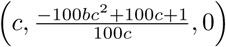 *of the system Eq.(2) is locally asymptotically stable if*

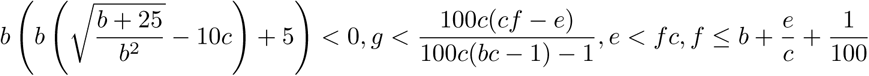

*Proof.* The fixed point 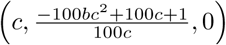 of the system Eq.(2) is locally asymptotically stable if all entries in the sequence {*a*_3_, *a*_2_, *H, a*_0_} where *H* = *a*_2_*a*_1_ − *a*_3_*a*_0_ are nonzero and of the same sign.

Here we see *a*_3_ = −1 which is negative and hence in order to have local stability of the fixed point, the coefficients *a*_2_, *a*_0_ and *H* are required to be negative. It can seen that the coefficients *a*_2_, *a*_0_ and *H* are negative provided

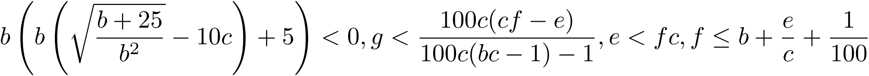

which can be seen by simple algebraic simplification.

Here we illustrate some example of local stability of the fixed point 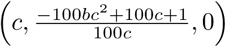 of the system Eq.(2). For *b* = 31, *c* = 0.391, and for all other positive parameters, the fixed point becomes(0.391, 0.0452, 0) which is locally asymptotically as shown in Fig.2.

**Figure 2:**
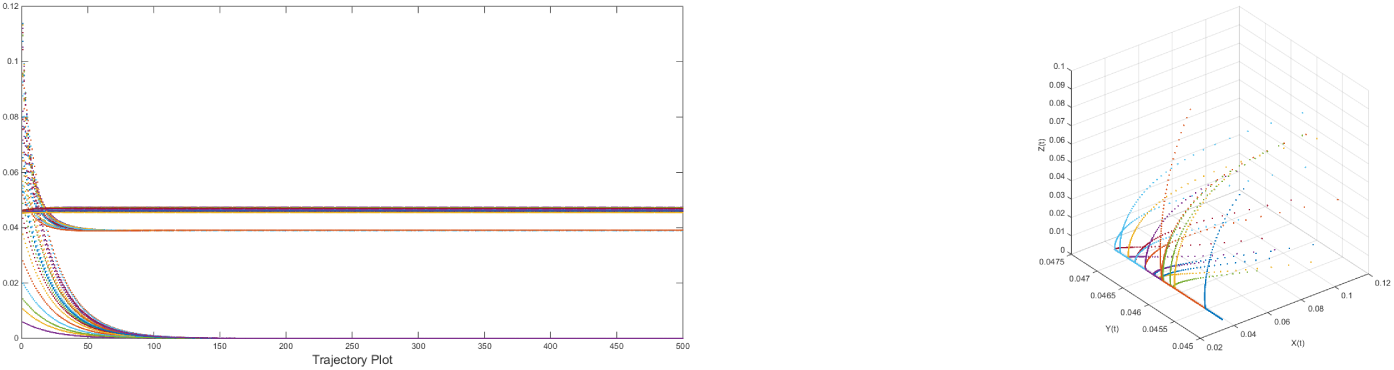
Left: Locally asymptotically stable trajectory of the fixed point(0.391, 0.0452, 0) for twenty different initial values(20 different set of colors are presented in the figure) taken from the neighbourhood of the fixed point, Right: corresponding three dimensional phase space.

The Jacobian evaluated at the fixed point 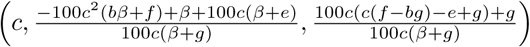 is

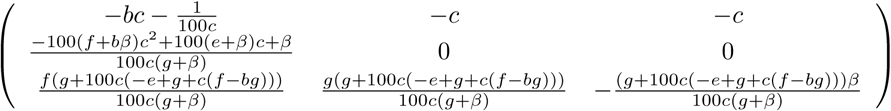

The coefficients of the characteristic polynomial *a*_3_, *a*_2_, *a*_1_ and *a*_0_ are given as:

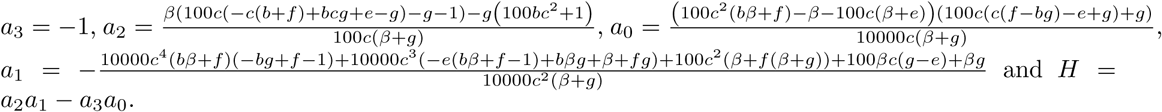

#### Theorem 2.4.

*The fixed point* 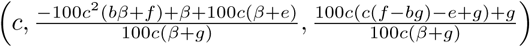 *of the system Eq.(2) is locally asymptotically stable if*

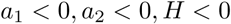

Here we illustrate the local behaviour of the fixed pint through an example.

We set the parameters *b* → 406; 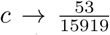, *e* → 6, 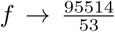 *g* → 36, *β* → 67 then the fixed point becomes(0.00332935, 1.725, 0.926866). This fixed point is locally asymptotically stable since all the eigenvalues(−62.0036, −4.44973, −0.00198725) are negative. The trajectories are shown n Fig. 3 for twenty different initial values taken from the neighbourhood of the fixed point.

**Figure 3:**
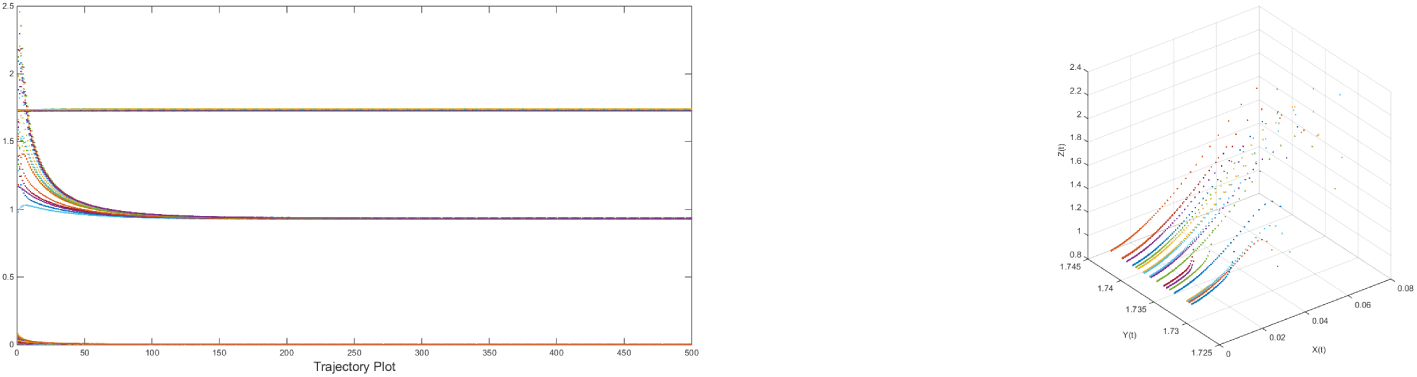
Left: Locally asymptotically stable trajectory of the fixed point(0.00332935, 1.725, 0.926866) for twenty different initial values(20 different set of colors are presented in the figure) taken from the neighbourhood of the fixed point, Right: corresponding three dimensional phase space.

For the fixed point 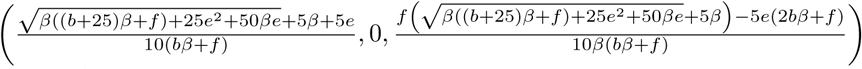 of the system stated in the Eq.(2), the Jacobian becomes

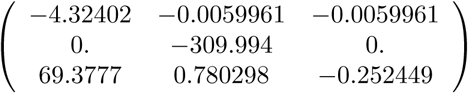

and its eigenvalues are −309.994, −4.21915 and −0.357321 when we consider the parameters *b* → 443, *c* → 310, *e* → 36 *f* → 6046, *g* → 68, *β→* 22. Hence the fixed point(0.0059961, 0, 0.011475) is locally asymptotically stable. The trajectory plots including its phase space for twenty different initial values which are taken from the neighbourhood of the fixed point are shown in Fig 4.

**Figure 4:**
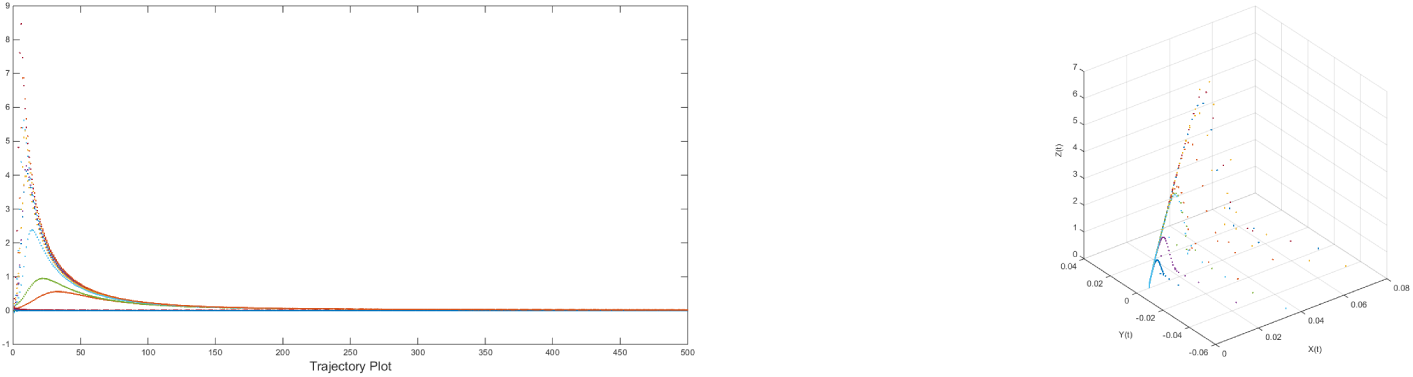
Left: Locally asymptotically stable trajectory of the fixed point(0.0059961, 0, 0.011475) for twenty different initial values(20 different set of colors are presented in the figure) taken from the neighbourhood of the fixed point, Right: corresponding three dimensional phase space.

### 2.2 Other Possible Behaviour of the System in the Case *I*11

The system Eq.(2) in the Case *I*11 also exhibits other kind of dynamical behavior such as high period, limit cycles and chaos(sensitive to the initial conditions). Some examples are shown in this section. The chaotic trajectory are confirmed through fractal(box counting dimension) of the three dimensional trajectory plots.

**Table 1:** Dynamics of System in the Case *I*11. The trajectory plots in the left side and the corresponding phase space are given in the right side of the are shown in the Fig. 5(Sl. 1to 6) for all the examples adumbrated in the Table 1.

| Serial No | Parameter: $(b, c, e, f, g, \beta)$ | Behaviour |
| --- | --- | --- |
| 1 | (0.0185, 0.7015, 0.9521, 0.7490, 0.7567, 0.5421) | Quasi-periodic |
| 2 | (0.5471, 0.1386, 0.1493, 0.2575, 0.8407, 0.2543) | Quasi-periodic |
| 3 | (0.2290, 0.6419, 0.4845, 0.1518, 0.7819, 0.1006) | Chaotic attractor (sensitive to the initial condition). Fractal (box counting) dimension for three dimensional trajectory is (1.145, 1.054, 1.287). |
| 4 | (0.1738, 0.6514, 0.4987, 0.2845, 0.8306, 0.8184) | Chaotic attractor, (sensitive to the initial condition). Fractal (box counting) dimension for three dimensional trajectory is (0.9477, 1.124, 1.035). |
| 5 | (0.0752, 0.5891, 0.6742, 0.5880, 0.5337, 0.4726) | Chaotic attractor (sensitive to the initial condition). Fractal (box counting) dimension for three dimensional trajectory is (0.6988, 1.124, 1.721). |
| 6 | (0.0966, 0.2654, 0.7919, 0.9369, 0.5683, 0.9380) | Attracting. The trajectories are converging to the fixed point (0.2653, 0.9907, 0.0212). |

**Figure 5:**
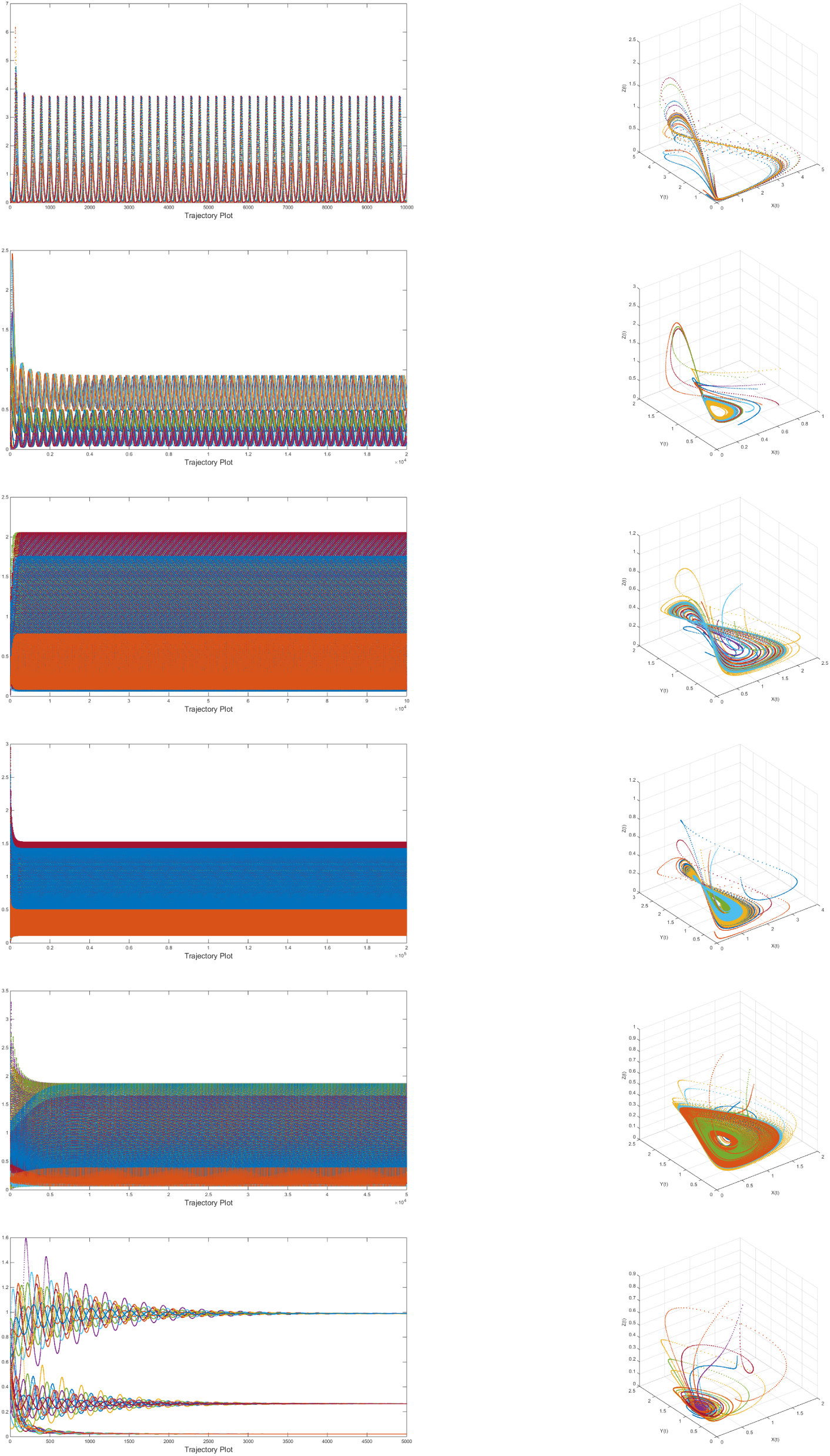
Left: Trajectory plot for twenty different initial values(20 different set of colors are presented in the figure), Right: corresponding three dimensional phase space.

In this case, we have seen there are four non-negative fixed points of the system Eq.(2). It is seen that for the fixed point 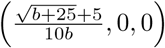 only prey persists and predator as well as omnivore both extinct. It is noted that the persistence of the prey depends on carrying capacity of the prey(b). For the fixed point 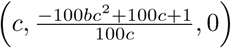, prey and predator coexist but the omnivore will extinct and this coexistence depends on death rate of the predator in the absence of the prey. For the fixed point 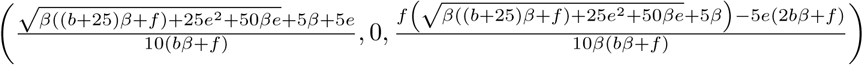 only prey and omnivore will coexist and that depends on all the parameters concerned in the system. The only other fixed point retains the coexistence of all the three species and that too depends on all the parameters involved in the system.

### 2.3 Stability of the Fixed Points of the System in the Case *I*12

The system Eq.(2) in the Case *I*12 has three non-negative fixed points 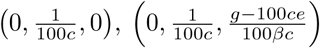, 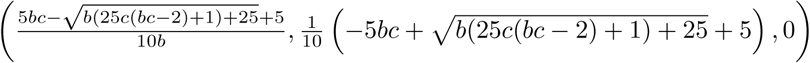.

Here the Jacobian evaluated at a fixed point is same as in the previous case. The immigration function(q=0.01) does not effect the partial derivatives.

#### Theorem 2.5.

*The fixed point* 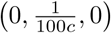 *of the system Eq.(2) is locally asymptotically stable if* 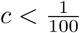 *and g* < 100*ce*

*Proof.* The Jacobian about the fixed point 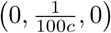 is

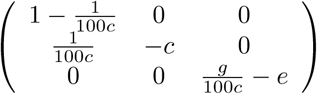

The eigenvalues are –*c*, 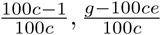. It can be easily deducible by doing simple algebraic simplification that all the real eigenvalues are negative if 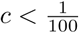 and *g <* 100*ce*.

Here we go with some example for illustration purpose of the local stability of the fixed point 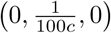.

For the parameters *c* → 0.00061875, *e* → 98., *g* → 2.70588 and with other non-negative parameters, the fixed point.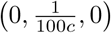 becomes(0, 16.1616, 0) which is attracting for twenty different initial values taken from very close neighbourhood of the fixed point The trajectory plot including its phase space are shown in Fig. 6(up). For the same parameters, we took initial values from a perturbed neighbourhood of fixed point, immediately then the fixed point repels as shown in Fig. 6(down). This repelling example triggers the chaotic solutions as the fixed point repels because of perturbed neighbourhood although the condition of asymptotic stability is confirmed.

**Figure 6:**
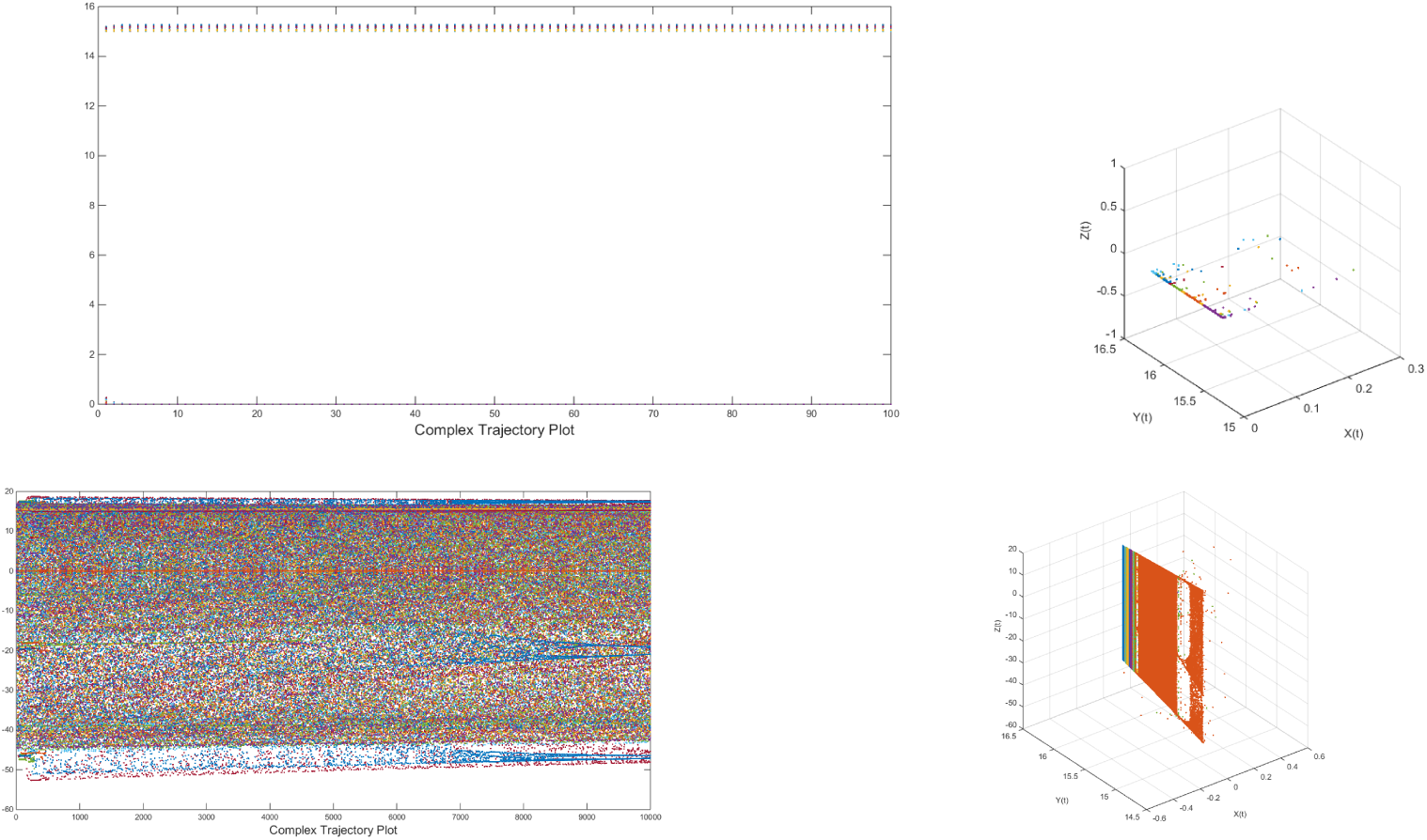
up left: Locally asymptotically stable trajectory of the fixed point(0, 16.1616, 0) for twenty different initial values(20 different set of colors are presented in the figure) taken from the neighbourhood of the fixed point, up Right: corresponding three dimensional phase space. down left: repelling trajectory of the fixed point(0, 16.1616, 0) and its corresponding phase space in the right down.

#### Theorem 2.6.

*The fixed point* 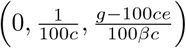 *of the system Eq.(2) is locally asymptotically stable if* 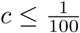. *and g >* 100*ce*.

*Proof.* Proof is left to the reader.

Some example for illustration purpose of the local stability of the fixed point 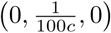 are given.

We set now the parameters 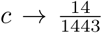, *e* → 17, *g* → 92, *β* → 31(which satisfy the Theorem 2.6), the fixed point becomes(0, 1.0307, 2.5105) which is attracting since the eigenvalues of the Jacobian are −0.00970201, −77.8257, −2.54122. The trajectory plot including its phase space is given in Fig. 7.

**Figure 7:**
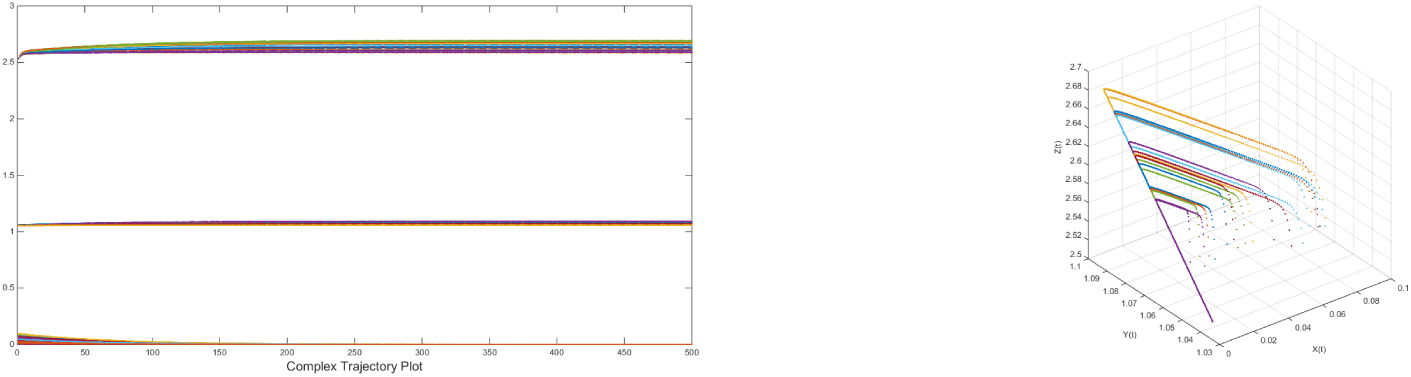
up left: Locally asymptotically stable trajectory of the fixed point(0, 1.0307, 2.5105) for twenty different initial values(20 different set of colors are presented in the figure) taken from the neighbourhood of the fixed point, up Right: corresponding three dimensional phase space.

Finding the condition of local asymptotic stability is just a matter of length calculations. We leave it. The jacobian of the fixed point 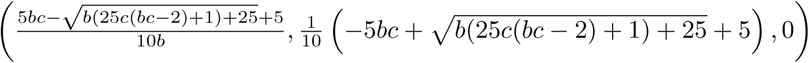 is very complicated. So we took specific example to adumbrate the attracting behaviour of the fixed point. We set the parameters 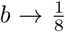, 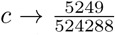, *g* → 4, *e* → 6, *f* → 1 with other non-negative parameters and then the fixed point becomes(0.0000116584, 0.9999, 0) which is attracting as shown in the Fig. 8 with its phase space.

**Figure 8:**
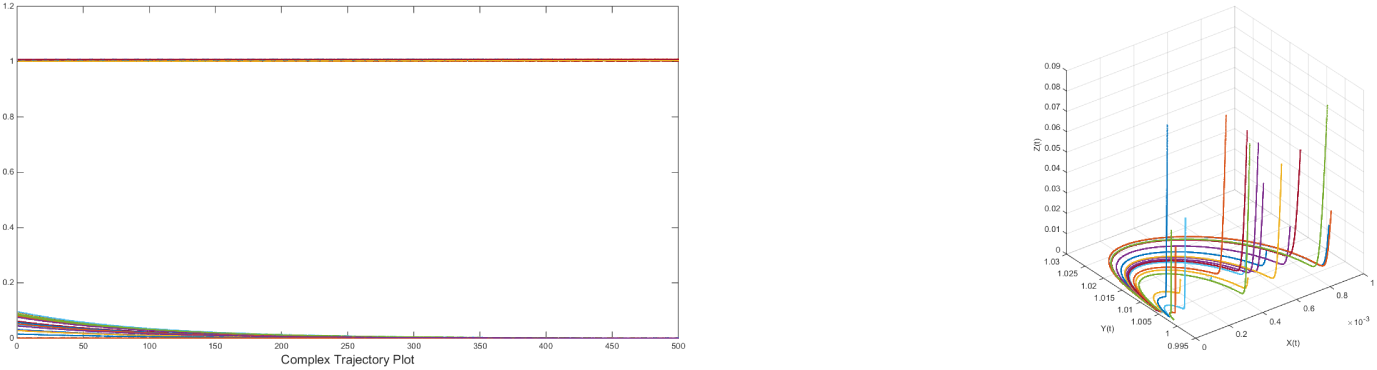
up left: Locally asymptotically stable trajectory of the fixed point(0.0000116584, 0.9999, 0) for twenty different initial values(20 different set of colors are presented in the figure) taken from the neighbourhood of the fixed point, up Right: corresponding three dimensional phase space.

### 2.4 Other Possible Behaviour of the System in the Case *I*12

In the Case *I*12 of the system Eq.(2), other kind of dynamical behavior such as high period, limit cycles and chaos(sensitive to the initial conditions) are seen. Some examples are shown in this section.

The trajectory plots in the left side and the corresponding phase space are given in the right side of the are shown in the Fig. 9(Sl. 1to 6) for all the examples adumbrated in the Table 2.

**Table 2:** Dynamics of System in the Case *I*12.

| Serial No | Parameter: $(b, c, e, f, g, \beta)$ | Behaviour |
| --- | --- | --- |
| 1 | $(0.3398, 0.2873, 0.1709, 0.3993, 0.6976, 0.2037)$ | Chaotic attractor, Fractal (box counting) dimension for three dimensional trajectory is $(0.854, 0.112, 0.784)$. |
| 2 | $(0.0954, 0.3526, 0.5934, 0.5852, 0.6677, 0.6480)$ | Chaotic attractor, Fractal (box counting) dimension for three dimensional trajectory is $(1.104, 1.002, 0.948)$. |
| 3 | $(0.9580, 0.0954, 0.0356, 0.8862, 0.2469, 0.0089)$ | Slowly attracting towards the fixed point $(0, 0.1235, 0.8627)$ |
| 4 | $(0.1397, 0.3939, 0.9806, 0.6448, 0.8964, 0.4822)$ | Quasi-periodic |
| 5 | $(0.8713, 0.0832, 0.4617, 0.0304, 0.7532, 0.7000)$ | Quasi-periodic |
| 6 | $(0.0966, 0.2654, 0.7919, 0.9369, 0.5683, 0.9380)$ | Attracting. The trajectories are converging to the fixed point $(0.2653, 0.9907, 0.0212)$. |

**Figure 9:**
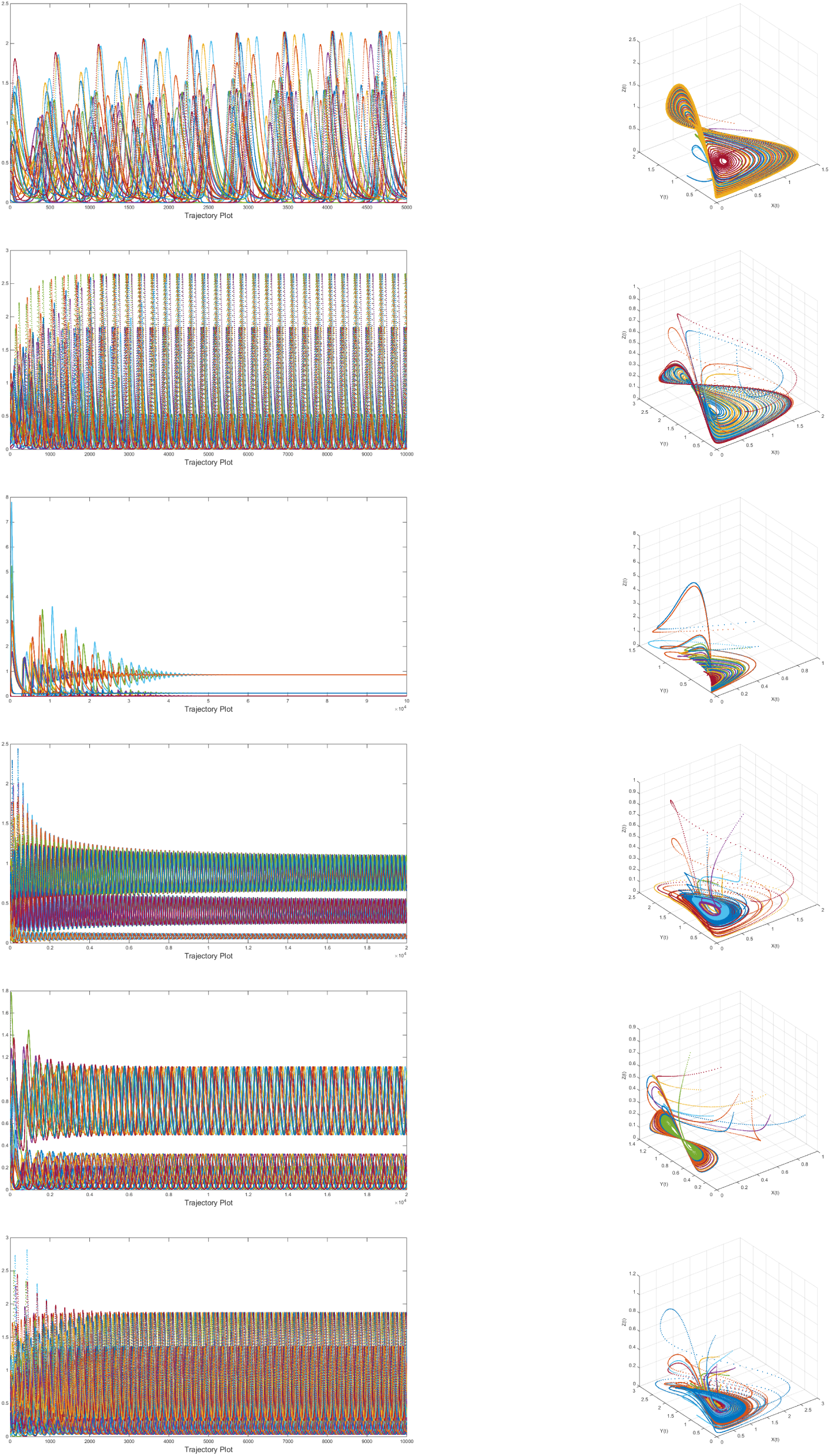
Left: Trajectory plot for twenty different initial values(20 different set of colors are presented in the figure), Right: corresponding three dimensional phase space.

Due to the effect of the small immigration(*q* = 0.01*i.e.*1%) of the predator in the system as in Case *I*12, there are three non-negative fixed points exist and none of which exhibits the coexistence of all the three species(prey, predator and omnivore). It is observed that only predator exists and that depends on the death rate of the predator in the absence of the prey(c) for the fixed point 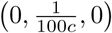. In the other fixed point 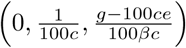, only predator and omnivore coexist and the prey population dies. For the rest fixed point, only prey and predator coexist and that coexistence depends on the parameter b and c i.e. carrying capacity of the prey and death rate of the predator in the absence of the prey.

### 2.5 Stability of the Fixed Points of the System in the Case *I*13

The system Eq.(2) in the Case *I*13 has three non-negative fixed points 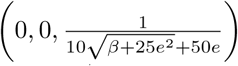, 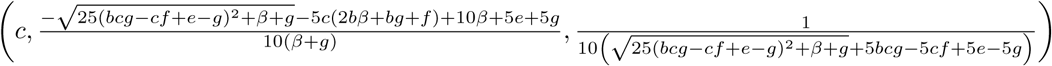 and 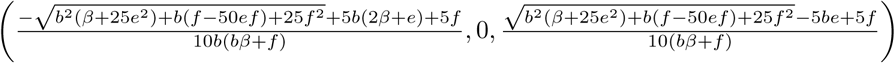.

#### Theorem 2.7.

*The fixed point* 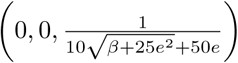 *of the system Eq.(2) is locally asymptotically stable if* 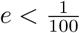 *and* 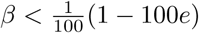

*Proof.* Proof is left to the reader.

Here by considering the parameters *c* → 188, 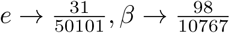 and other non-negative parameters, the fixed point 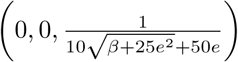 becomes(0, 0, 1.01474) which is locally asymptotically stable because the eigenvalues(−188, −0.014737, −0.0190908) of the corresponding Jacobian are all negative. The trajectory plot for twenty different initial values taken from the very close neighbourhood of the fixed point, including the phase space are given in the Fig. 10.

**Figure 10:**
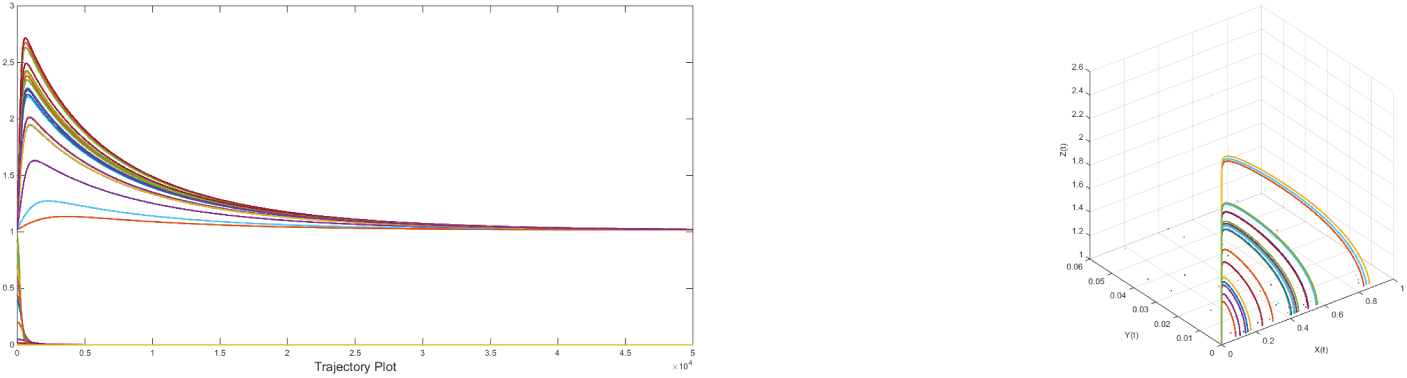
Left: Locally asymptotically stable trajectory of the fixed point(0, 0, 1.01474) for twenty different initial values(20 different set of colors are presented in the figure) taken from the neighbourhood of the fixed point, Right: corresponding three dimensional phase space.

The Jacobian of the system with respect to the fixed point 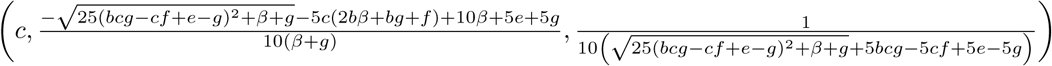 is very much complicated and hence finding the condition for local asymptotic stability is tedious. So we prefer to choose some specific parameters in order to see the local stability of the fixed point. Here we go with an example.

We consider the parameters *b* → 335, 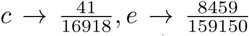, *f* → 38, *g* → 76, *β* → 51 and then the fixed point becomes(0.00242345, 0.0745536, 0.113589) which is locally asymptotically stable because the eigenvalues(−5.87907, −0.813321, −0.000548423) are all negative. The trajectory plot including its phase space are given in the following Fig. 11.

**Figure 11:**
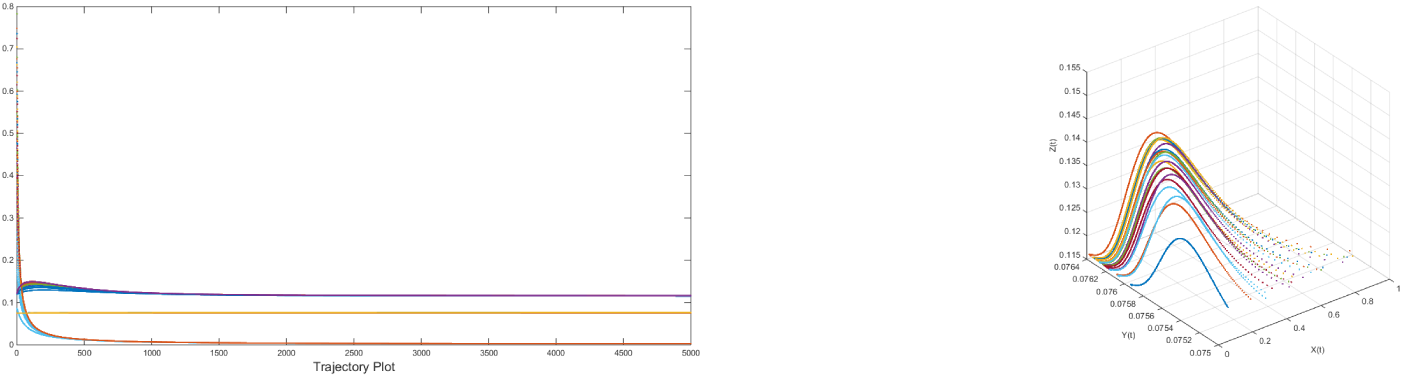
Left: Locally asymptotically stable trajectory of the fixed point(0.00242345, 0.0745536, 0.113589) for twenty different initial values(20 different set of colors are presented in the figure) taken from the neighbourhood of the fixed point, Right: corresponding three dimensional phase space.

Again, the Jacobian of the system with respect to the fixed point 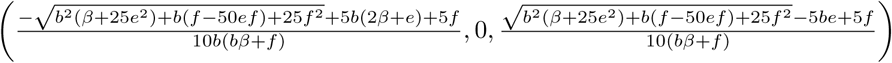 is very much complicated and hence finding the condition for local asymptotic stability is tedious. So we prefer to choose specific parameters in understanding the local stability of the fixed point. Here we go with the following example.

We consider the parameters *b*→52, *c* → 415, *e* → 87, *f* → 74, *g* → 31, *β* → 96 and then the fixed point becomes(0.0192285, 0, 0.000116838) which is locally asymptotically stable because the eigenvalues(−414.981, −85.5995, −0.999885) are all negative. The trajectory plot including its phase space are given in the following Fig. 12.

**Figure 12:**
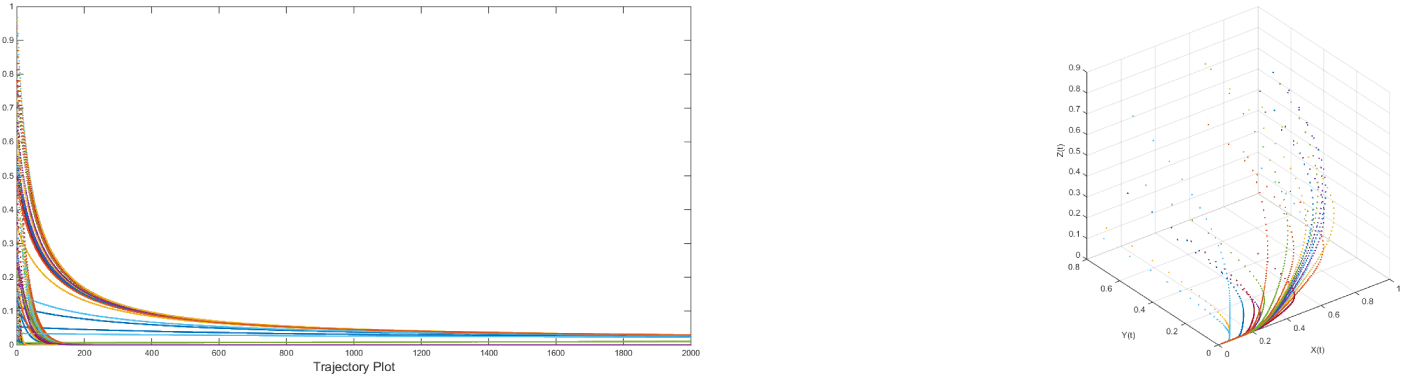
Left: Locally asymptotically stable trajectory of the fixed point(0.0192285, 0, 0.000116838) for twenty different initial values(20 different set of colors are presented in the figure) taken from the neighbourhood of the fixed point, Right: corresponding three dimensional phase space.

### 2.6 Other Possible Behaviour of the System in the Case *I*13

in the Case *I*13 of the system Eq.(2), other kind of dynamical behavior such as high period, limit cycles and chaos(sensitive to the initial conditions) are seen. Some examples are shown in this section.

**Figure 13:**
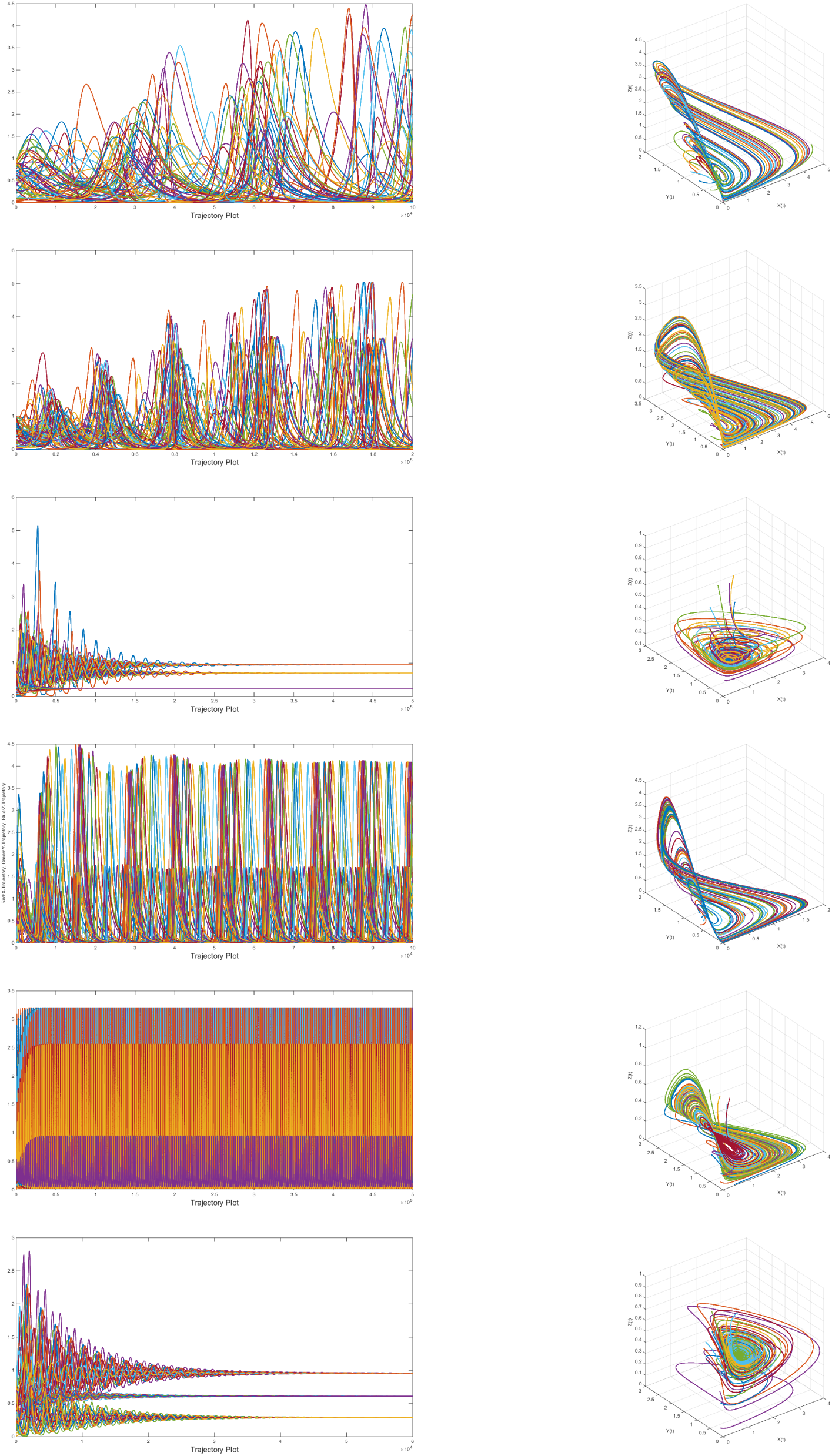
Left: Trajectory plot for twenty different initial values(20 different set of colors are presented in the figure), Right: corresponding three dimensional phase space.

**Table 3:** Dynamics of System in the Case *I*131. The trajectory plots in the left side and the corresponding phase space are given in the right side of the are shown in the Fig. 13 for all the examples adumbrated in the Table 1.

| Serial No | Parameter: $(b, c, e, f, g, \beta)$ | Behaviour |
| --- | --- | --- |
| 1 | (0.0350, 0.4055, 0.2497, 0.4809, 0.8808, 0.2807) | Chaotic attractor, Fractal (box counting) dimension for three dimensional trajectory is (1.105, 1.145, 0.983). |
| 2 | (0.1230, 0.5800, 0.3285, 0.2682, 0.5502, 0.1805) | Chaotic attractor, Fractal (box counting) dimension for three dimensional trajectory is (0.7384, 0.587, 0.879). |
| 3 | (0.0828, 0.9504, 0.0164, 0.1147, 0.0124, 0.2162) | Slowly attracting towards the fixed point (0.9504, 0.6989, 0.2225) |
| 4 | (0.4003, 0.1119, 0.4243, 0.6135, 0.9881, 0.2199) | Very high period (forming large limit cycle) |
| 5 | (0.1589, 0.8109, 0.4765, 0.1163, 0.8757, 0.6352) | Very high period |
| 6 | (0.1024, 0.9591, 0.1529, 0.1525, 0.1556, 0.0896) | Slowly attracting. The trajectories are converging to the fixed point (0.9572, 0.2905, 0.6121). |

Small immigration(*r* = 0.01*i.e.*1%) of the omnivore in the system as in Case *I*13 leading to different dynamical behavior. There are three non-negative fixed points exist and one of which exhibits the co-existence of all the three species(prey, predator and omnivore). It is observed that only omnivore exists for the fixed point 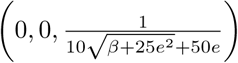; and that existence of omnivore depends on the death rate of the omnivo in the absence of its food prey and predator. On the other side, for the other fixed point 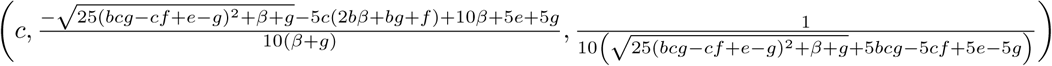, 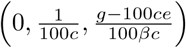 only prey and omnivore coexist. For the rest fixed point, all three species exist that coexistence depends on all the parameters involved in the system.

### 2.7 Stability of the Fixed Points of the System in the Case *J* 11

In this case, system has changed due to effect a certain percentage with respect to the number of prey of the population, i.e. here *p*(*x*) = *px*. Accordingly the system Eq.(2) has five non-negative fixed points 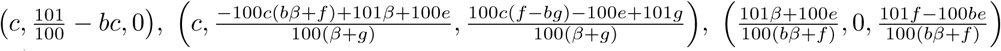 and 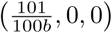 and (0, 0, 0).

The Jacobian of the system according would be at a fixed point 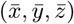:

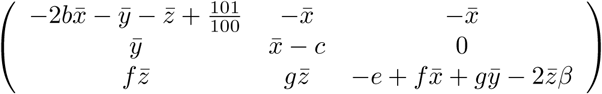

#### Theorem 2.8.

*The fixed point* 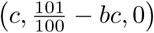 *is locally asymptotically stable if,*

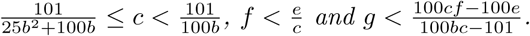

*Proof.* Proof is left to the reader.

We shall here exhibit numerical example in order to see the local stability of the fixed point 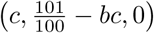. We consider the parameters *b* → 267,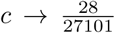, *e* → 53, *f* → 54, *g* → 21, *β* → 31 then the fixed point becomes(0.00103317, 0.734143, 0) which is attracting(all the eigenvalues are negative)) as shown in Fig. 14 with its trajectory plot and phase space.

**Figure 14:**
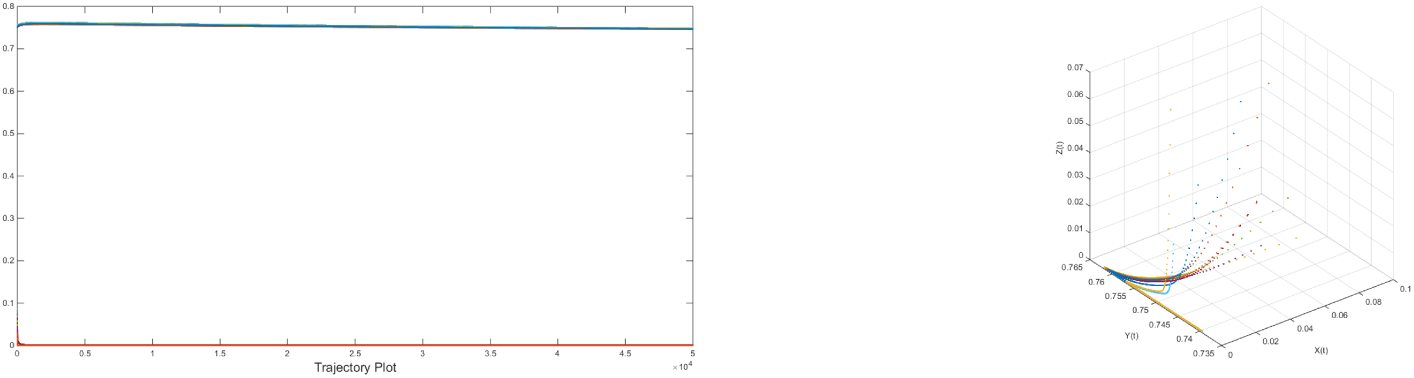
Left: Attracting trajectory of the fixed point(0.00103317, 0.734143, 0) for twenty different initial values(20 different set of colors are presented in the figure) taken from the neighbourhood of the fixed point, Right: corresponding three dimensional phase space.

The Jacobian corresponding to the fixed point 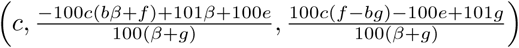 is bit complicated and hence deducing the condition for local stability is tedious. Hence we are just describing the stability of the fixed point through an example illustrated below.

We consider *b* → 167, 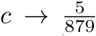, *e* → 81, 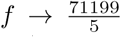, *g* → 83, *β* → 20 then the fixed point becomes (0.0057, 0.0116615, 0.0483954) which is locally asymptotically stable as the real part of the eigenvalues(−0.958891 + 1.97988*i,* −0.958891 −1.97988*i,* −0.0000683258) of the Jacobian are negative. The trajectory plot including its phase space are given in Fig. 15.

**Figure 15:**
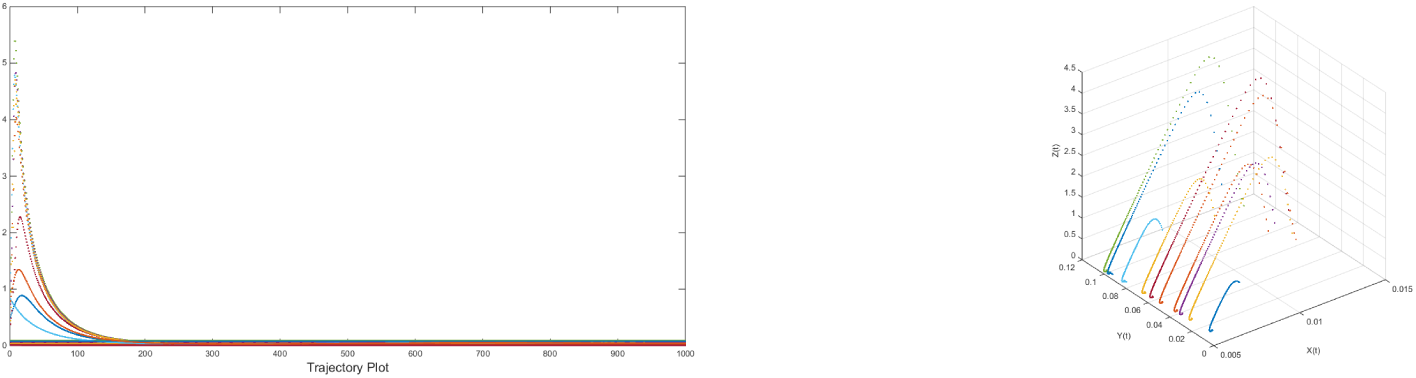
Left: Attracting trajectory of the fixed point(, 0.0116615, 0.0483954) for twenty different initial values(20 different set of colors are presented in the figure) taken from the neighbourhood of the fixed point, Right: corresponding three dimensional phase space.

#### Theorem 2.9.

*The fixed point* 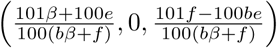 *is locally asymptotically stable if* 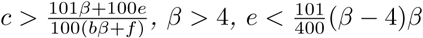 *and* 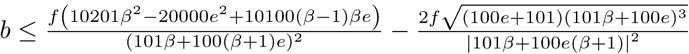

*Proof.* Proof is left to the reader.

Here we present one example for illustration of the local stability of the fixed point 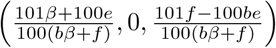.

We take the parameters {*b* → 339, *c* → 273, *e* → 14, *f* → 4782, *g* → 31, *β* → 43 } and then the fixed point becomes(0.00432977, 0, 0.00296658) which is attracting since the eigenvalues(−272.997, −0.92222, −0.269631) of the Jacobian are all negative. The trajectory plots as well as the phase space are given in Fig. 16.

**Figure 16:**
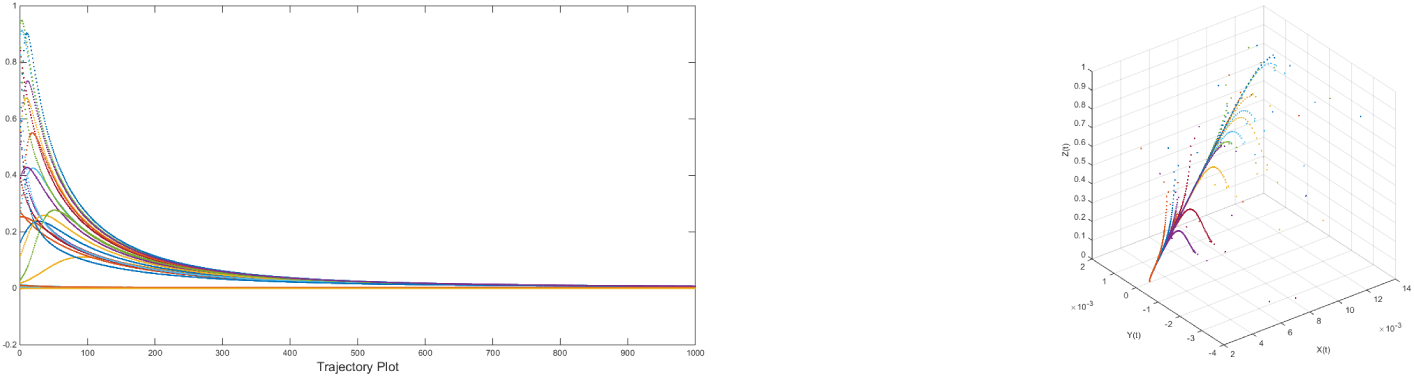
Left: Attracting trajectory of the fixed point(, 0.0116615, 0.0483954) for twenty different initial values(20 different set of colors are presented in the figure) taken from the neighbourhood of the fixed point, Right: corresponding three dimensional phase space.

#### Theorem 2.10.

*The fixed point* 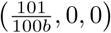 *is locally asymptotically stable if*

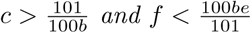

*Proof.* Proof is left to the reader.

We consider the parameters *b* → 31, *c* → 98, *e* → 46, *f* → 17 and other non-negative parameters, then the fixed point becomes(0.0325806, 0, 0) which is attracting as the parameters satisfy the Theorem 2.10. The trajectories and phase space are given in Fig. 17.

**Figure 17:**
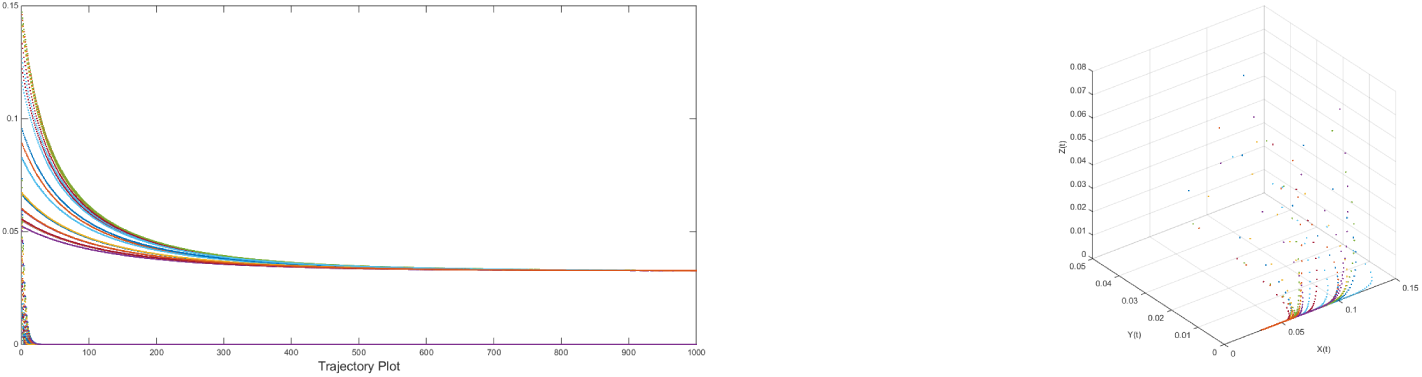
Left: Attracting trajectory of the fixed point(0.0327, 0, 0) for twenty different initial values(20 different set of colors are presented in the figure) taken from the neighbourhood of the fixed point, Right: corresponding three dimensional phase space.

Small immigration(*p*(*x*) = *px*) relative to the density of the prey in the system as in Case *J* 11 leading to different dynamical behavior. There are five non-negative fixed points exist and one of which is not locally asymptotically unstable. One of the fixed points 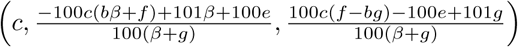 retains the coexistence of all the three species(prey, predator and omnivore) where all the parameters of the system involved. It is observed that only omnivore vanishes(prey and predator coexist) for the fixed point 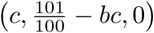 and that existence of prey and predator depend on the carrying capacity of the prey and on the death rate of the predator in the absence of prey. On the other side, for the other fixed point 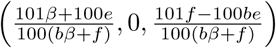 only prey and omnivore coexist. For the rest fixed point, only prey exists and that depends on the carrying capacity of the prey.

#### Theorem 2.11.

*The fixed point* (0, 0, 0) *is asymptotically unstable for any non-negative parameters involved in the system.*

*Proof.* One of the eigenvalues of the Jacobian corresponding to the fixed point origin(0, 0, 0) is 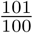 and consequently the origin cannot be locally asymptotically stable.

Here we put few examples of unstablity of the fixed point(0, 0, 0). Since the origin is unstable, the repelling trajectories yield to formation of chaotic and limit cycle attractors.

- (*b, c, e, f, g, β*) =(0.3968, 0.0740, 0.6841, 0.4024, 0.9828, 0.4022) then the origin repels and forms a limit cycle as seen in the phase space given in the following Fig. 18 from top(1).
- (*b, c, e, f, g, β*) =(0.0503, 0.2287, 0.8342, 0.0156, 0.8637, 0.0781) then the origin repels and forms a chaotic attractor as seen in the phase space given in the following Fig. 17 from top(2).
- (*b, c, e, f, g, β*) =(0.2462, 0.3427, 0.3757, 0.5466, 0.5619, 0.3958) then the origin repels and forms a limit cycle as seen in the phase space given in the following Fig. 18 from top(3).
- (*b, c, e, f, g, β*) =(0.0721, 0.4067, 0.6669, 0.9337, 0.8110, 0.4845) then the origin repels and forms a limit cycle as seen in the phase space given in the following Fig. 18 from top(4).
- (*b, c, e, f, g, β*) =(0.1375, 0.3900, 0.9274, 0.9175, 0.7136, 0.6183) then the origin repels and forms a limit cycle as seen in the phase space given in the following Fig. 18 from top(5).
- (*b, c, e, f, g, β*) =(0.9345, 0.1079, 0.1822, 0.0991, 0.4898, 0.1932) then the origin repels and forms a limit cycle as seen in the phase space given in the following Fig. 18 from top(6).

### 2.8 Stability of the Fixed Points of the System in the Case *J* 12

In this case, system has changed due to effect a certain percentage with respect to the number of predator of the population, i.e. here *p*(*x*) = *qy*. Accordingly the system Eq.(2) has five non-negative fixed points 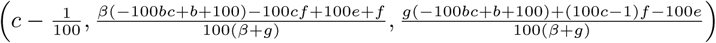,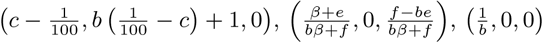 and (0, 0, 0).

**Figure 18:**
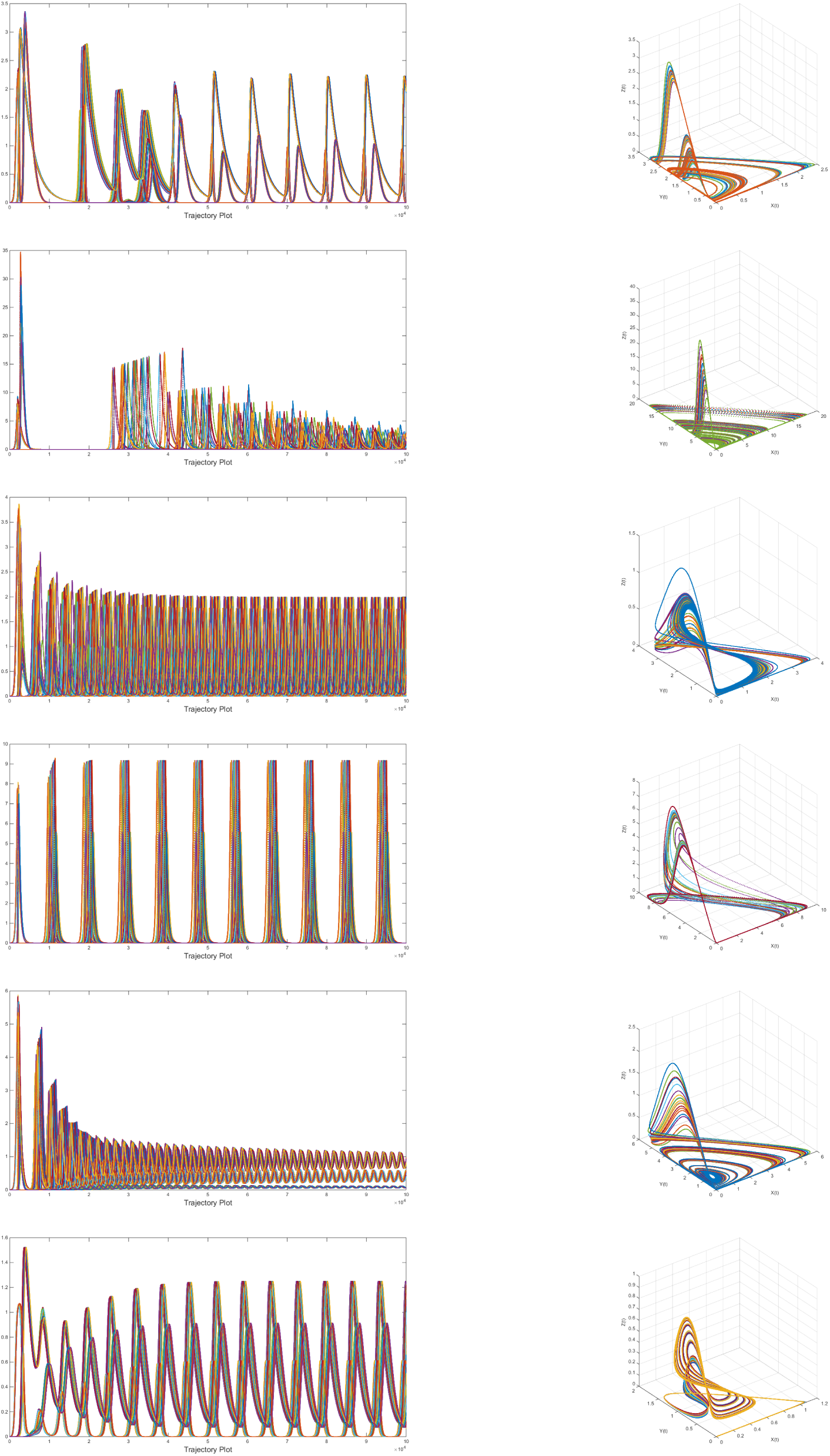
Left: Repelling trajectories from the fixed point(0, 0, 0) for twenty different initial values(20 different set of colors are presented in the figure) taken from the neighbourhood of the origin, Right: corresponding three dimensional phase space.

The Jacobian of the system according would be at a fixed point 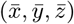:

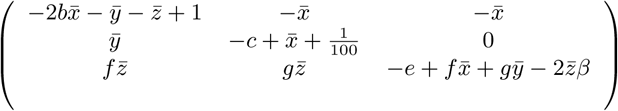

The fixed point 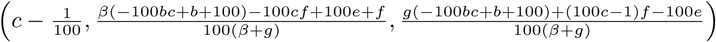 of the system in this case would be non-negative if and only if the following condition is satisfied.

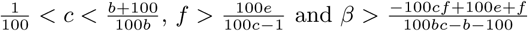

As the Jacobian corresponding to the fixed point is bit complicated so we proceed through an example to illustrate the local behavior. Consider the parameters *b* → 167, 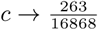, *e* → 81, 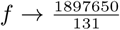, *g* → 83, *β* → 20 then the fixed point becomes(0.00559165, 0.0128532, 0.0533408) which is locally asymptotically stable since the real part of the eigenvalues(−1.00027 + 2.07754*i*, −1.00027 −2.07754*i*, −0.0000742684) of the Jacobian about the fixed point are all negative. The trajectory plot including the phase space are given in Fig. 19.

**Figure 19:**
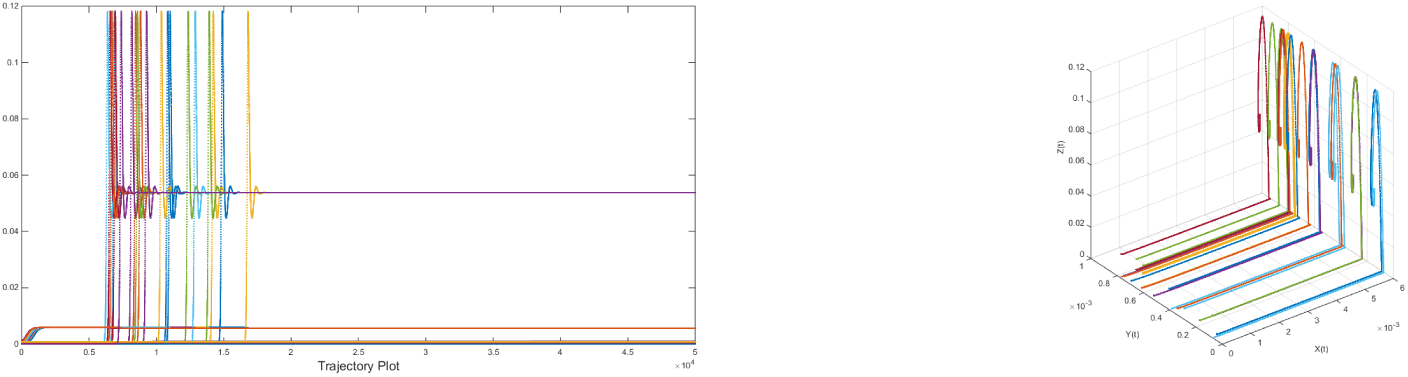
Left: Attracting trajectory of the fixed point(0.0057, 0.0004, 0.0538) for twenty different initial values(20 different set of colors are presented in the figure) taken from the neighbourhood of the fixed point, Right: corresponding three dimensional phase space.

#### Theorem 2.12.

*the fixed point* 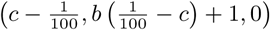 *is locally asymptotically stable if*

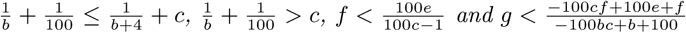

*Proof.* Proof is left to the reader.

Having set with the parameters *b* → 267,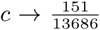, *e* → 53, *f* → 54, *g* → 21, *β* → 31, the fixed point becomes(0.0010, 0.7241, 0) which is attracting as the parameters satisfy the Theorem 2.12. The trajectory plot including its phase space are given in the Fig. 20.

**Figure 20:**
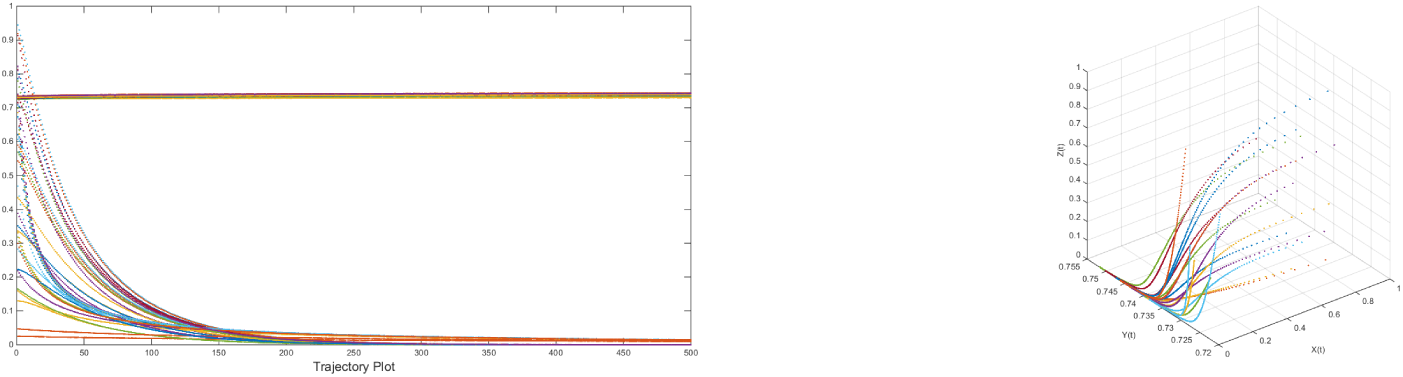
Left: Attracting trajectory of the fixed point(0.0010, 0.7381, 0) for twenty different initial values(20 different set of colors are presented in the figure) taken from the neighbourhood of the fixed point, Right: corresponding three dimensional phase space.

#### Theorem 2.13.

*the fixed point* 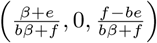 *is locally asymptotically stable if*

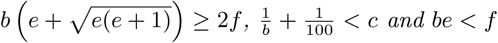

*Proof.* Proof is left to the reader.

Here we take the parameters *b* → 343, *c* → 88, *e* → 5, 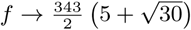, *g* → 31, *β* → 273 then the fixed point becomes(0.0029, 0, 0.0009) which is locally asymptotically stable as the parameters satisfy the Theorem 2.13. The trajectory plot including corresponding phase space are graphed in the following Fig. 21.

**Figure 21:**
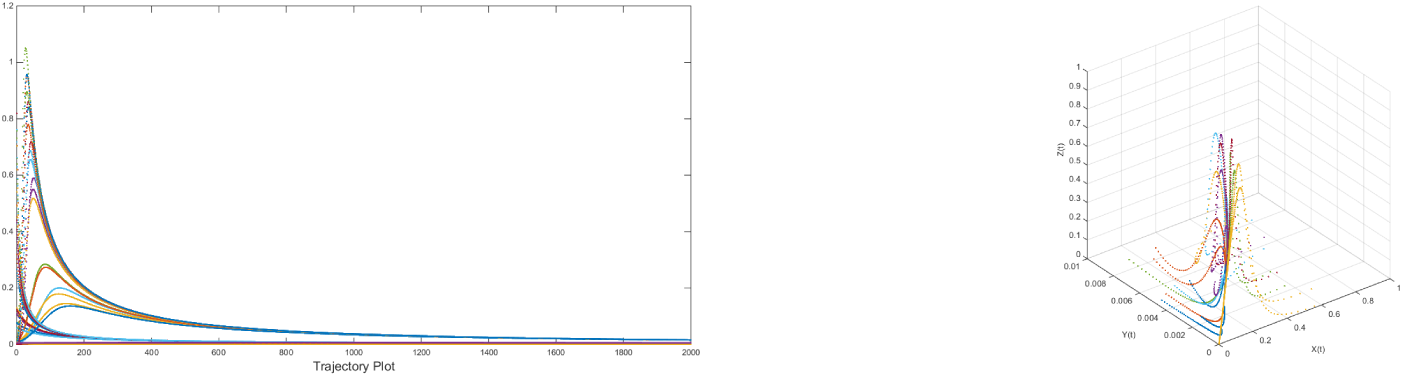
Left: Attracting trajectory of the fixed point(0.0029, 0, 0.0009) for twenty different initial values(20 different set of colors are presented in the figure) taken from the neighbourhood of the fixed point, Right: corresponding three dimensional phase space.

#### Theorem 2.14.

*the fixed point* 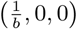 *is locally asymptotically stable if*

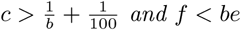

*Proof.* Proof is left to the reader.

Here we fix the parameters *b* → 170, *c* → 17, *e* → 52, *f* → 75, *g* → 29, *β* → 120 which satisfy the Theorem 2.14. So the fixed point becomes(0.00588235, 0, 0) which is locally asymptotically stable as it is seen in the Fig. 22.

**Figure 22:**
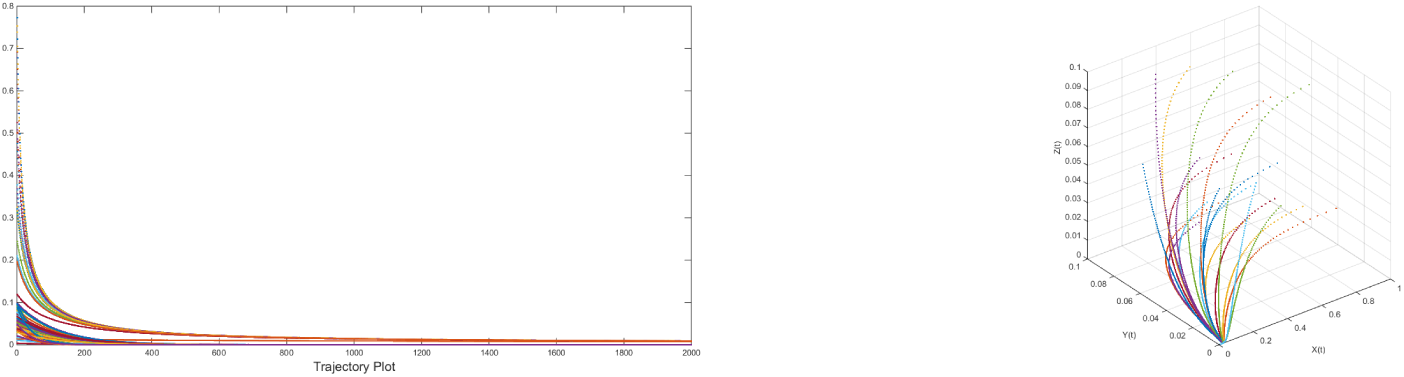
Left: Attracting trajectory of the fixed point(0.0059, 0, 0)for twenty different initial values(20 different set of colors are presented in the figure) taken from the neighbourhood of the fixed point, Right: corresponding three dimensional phase space.

Small immigration(*q*(*x*) = *qx*) relative to the density of the predator in the system as in Case J 12 leading to different dynamical behavior. There are five non-negative fixed points exist and one of which is not locally asymptotically unstable. One of the fixed points 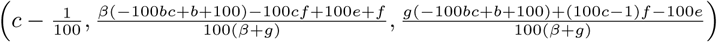 retains the coexistence of all the three species(prey, predator and omnivore) where all the parameters of the system involved. It is observed that only omnivore vanishes(prey and predator coexist) for the fixed point 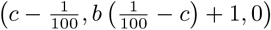 and that existence of prey and predator depend on the carrying capacity of the prey and on the death rate of the predator in the absence of prey. On the other side, for the other fixed point 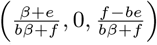 only prey and omnivore coexist. For the rest fixed point, only prey exists and that depends on the carrying capacity of the prey.

#### Theorem 2.15.

*The fixed point(*0, 0, 0) *is asymptotically unstable for any non-negative parameters involved in the system.*

*Proof.* One of the eigenvalues of the Jacobian corresponding to the fixed point origin(0, 0, 0) is 1 and consequently the origin cannot ne locally asymptotically stable.

Here we put few examples of unstablity of the fixed point(0, 0, 0). Since the origin is unstable, the repelling trajectories yield to formation of chaotic and limit cycle attractors.

- (*b, c, e, f, g, β*) =(0.0545, 0.5004, 0.4328, 0.9043, 0.6302, 0.9830) then trajectories are slowly convergent to the fixed point(0.4977, 0.5874, 0.3873) as shown in the following Fig. 23 from top(1).
- (*b, c, e, f, g, β*) =(0.1805, 0.6785, 0.0557, 0.0341, 0.2865, 0.0774) then the origin repels and forms a chaotic attractor as seen in the phase space given in the following Fig. 23 from top(2).
- (*b, c, e, f, g, β*) =(0.1528, 0.4057, 0.3125, 0.6939, 0.8907, 0.4907) then the origin repels and forms a chaotic attractor as seen in the phase space given in the following Fig. 23 from top(3).
- (*b, c, e, f, g, β*) =(0.0047, 0.6500, 0.6785, 0.2536, 0.8432, 0.2940) then the origin repels and forms a limit cycle as seen in the phase space given in the following Fig. 23 from top(4).
- (*b, c, e, f, g, β*) =(0.1275, 0.4962, 0.3105, 0.5786, 0.9436, 0.4269) then the origin repels and forms a chaotic attractor as seen in the phase space given in the following Fig. 23 from top(5).
- (*b, c, e, f, g, β*) =(0.1577, 0.6005, 0.9375, 0.1078, 0.9000, 0.5505) then trajectories are slowly convergent to the fixed point(0.5905, 0.9069, 0) as shown in the following Fig. 23 from top(6).

### 2.9 Stability of the Fixed Points of the System in the Case *J* 13

In this case, system has changed due to effect a certain percentage with respect to the number of omnivore of the population, i.e. here *r*(*x*) = *rz*. Accordingly the system Eq.(2) has five non-negative fixed points 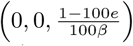,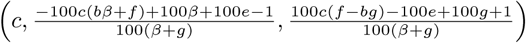, (*c*, 1 – *bc*, 0),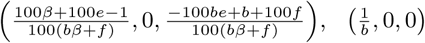 and(0, 0, 0).

**Figure 23:**
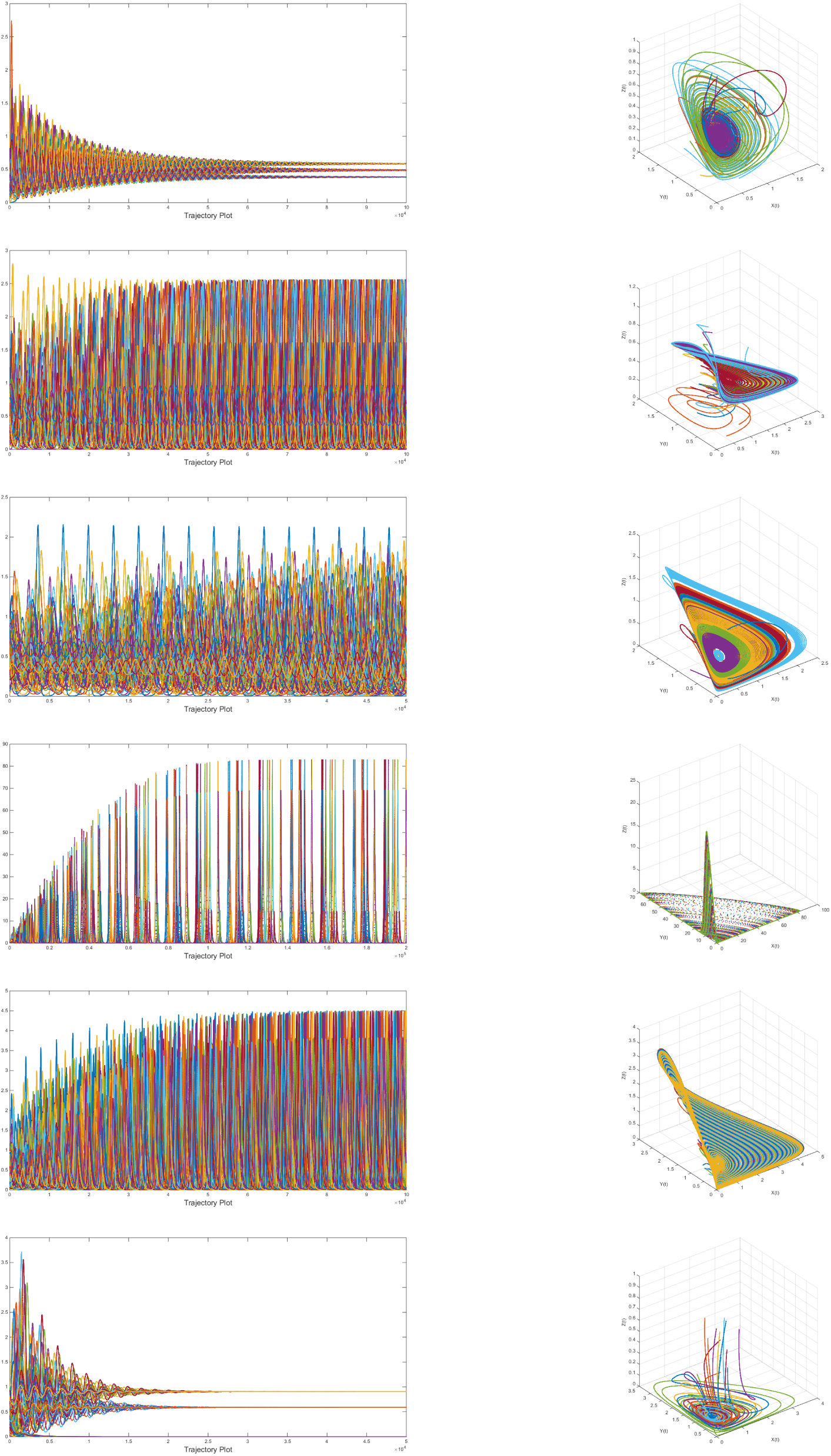
Left: Repelling trajectories from the fixed point(0, 0, 0) for twenty different initial values(20 different set of colors are presented in the figure) taken from the neighbourhood of the origin, Right: corresponding three dimensional phase space.

The Jacobian of the system according would be at a fixed point 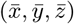:

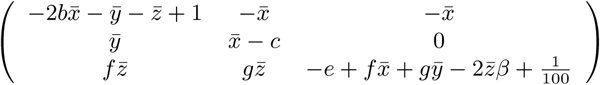

#### Theorem 2.16.

*The fixed point* 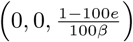 *is locally asymptotically stable if*

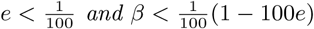

*Proof.* Proof is left to the reader.

Here we set the parameters 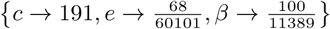 and then the fixed pint becomes (0, 0, 1.001) which is locally asymptotically stable since the parameters satisfy the Theorem 2.16. The trajectory plot and phase space are given in Fig. 24.

**Figure 24:**
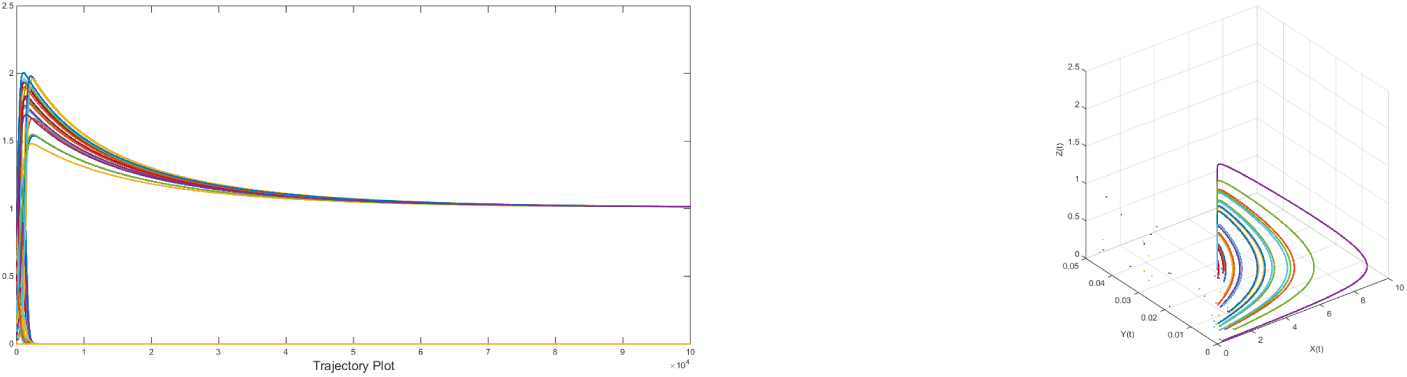
Left: Attracting trajectory of the fixed point(0, 0, 1.001) for twenty different initial values(20 different set of colors are presented in the figure) taken from the neighbourhood of the fixed point, Right: corresponding three dimensional phase space.

Here we discuss the local stability of the fixed point 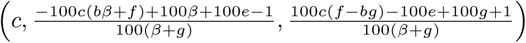 through an example. Here we fix the parameters *b* → 343, 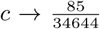, *e* → 82, *f* → 33499, *g* → 1, *β* → 53 and then the fixed point becomes(0.00245353, 0.151789, 0.00665095) which is locally asymptotically stable since the real part of the eigenvalues of the corresponding Jacobian are negative. The trajectory plot and phase space are given in Fig. 25.

**Figure 25:**
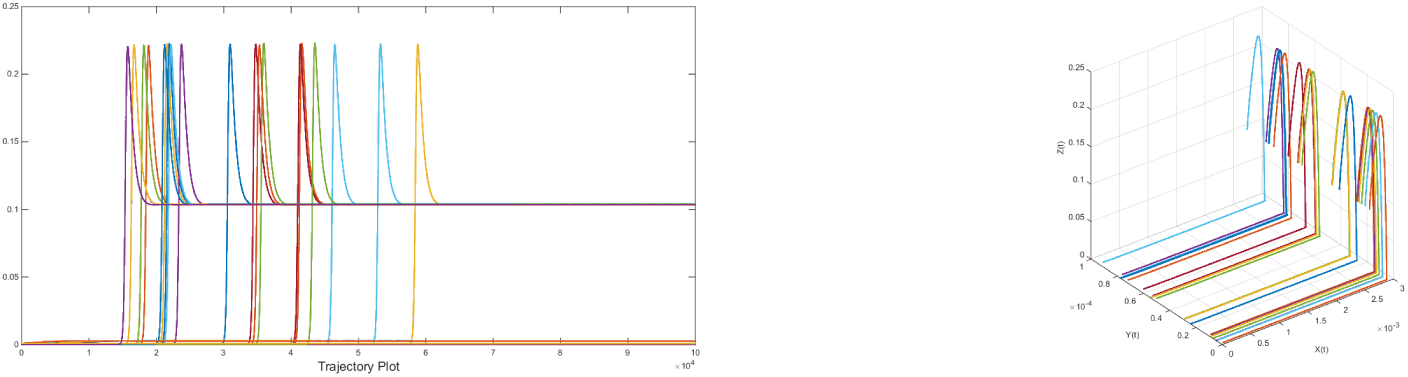
Left: Attracting trajectory of the fixed point(0.002, 0.16, 0.007) for twenty different initial values(20 different set of colors are presented in the figure) taken from the neighbourhood of the fixed point, Right: corresponding three dimensional phase space.

#### Theorem 2.17.

*The fixed point (c,* 1 − *bc*, 0) *is locally asymptotically stable if*

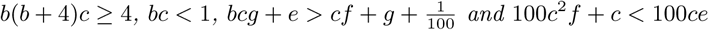

*Proof.* Proof is left to the reader.

Here we set the parameters *b* → 267, 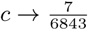, *e* → 53, *f* → 54, *g* → 21, *β* → 31 and then the fixed pint becomes(0.0010, 0.7269, 0) which is locally asymptotically stable since the parameters satisfy the Theorem 2.17. The trajectory plot and phase space are given in Fig. 26.

**Figure 26:**
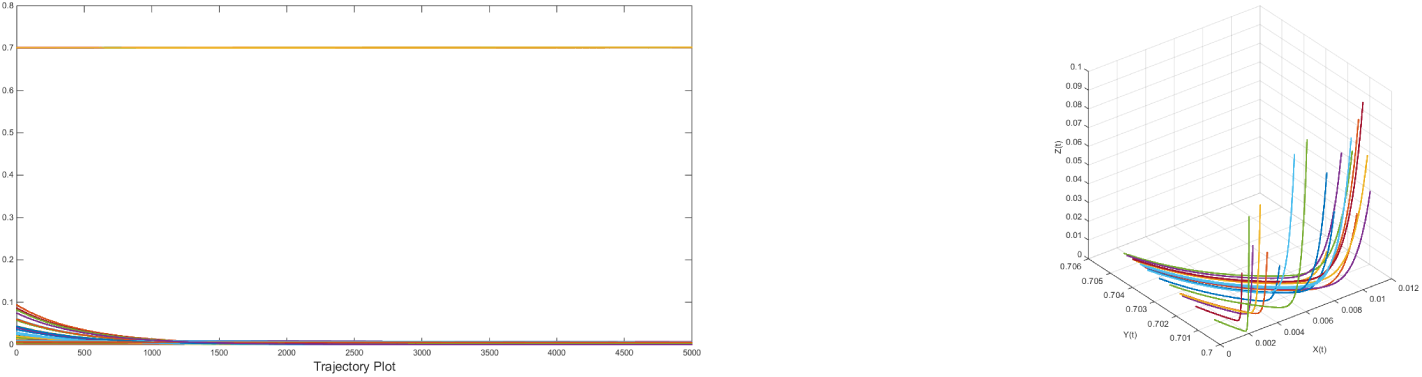
Left: Attracting trajectory of the fixed point(0.0010, 0.7269, 0) for twenty different initial values(20 different set of colors are presented in the figure) taken from the neighbourhood of the fixed point, Right: corresponding three dimensional phase space.

#### Theorem 2.18.

*The fixed point* 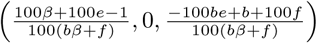 *is locally asymptotically stable if*

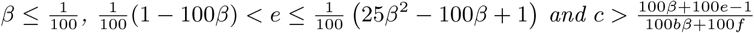

*Proof.* Proof is left to the reader.

Here we set the parameters *b* → 52, *c* → 415, *e* → 87, *f* → 4597, *g* → 31, *β* → 96 and then the fixed pint becomes(0.0190833, 0, 0.00766712) which is locally asymptotically stable since the parameters satisfy the Theorem 2.18. The trajectory plot and phase space are given in Fig. 27.

**Figure 27:**
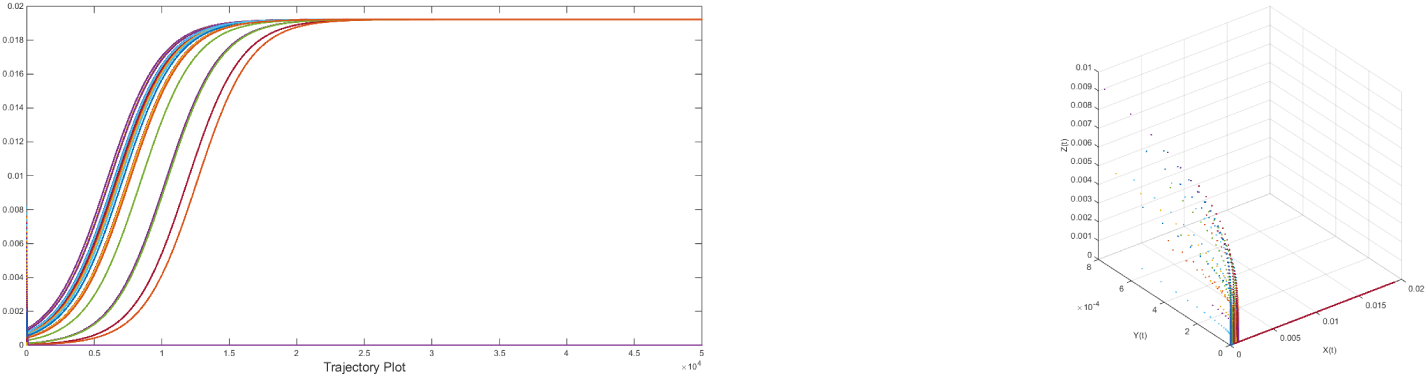
Left: Attracting trajectory of the fixed point(0.0191, 0, 0.0077) for twenty different initial values(20 different set of colors are presented in the figure) taken from the neighbourhood of the fixed point, Right: corresponding three dimensional phase space.

#### Theorem 2.19.

*the fixed point* 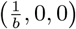 *is locally asymptotically stable if*

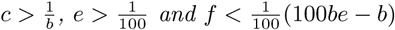

*Proof.* Proof is left to the reader.

Here we fix the parameters *b* → 68, *c* → 100, *e* → 95, *f* → 63 with other non-negative parameter which satisfy the Theorem 2.19. So the fixed point becomes(0.0147, 0, 0) which is locally asymptotically stable as it is seen in the Fig. 28.

**Figure 28:**
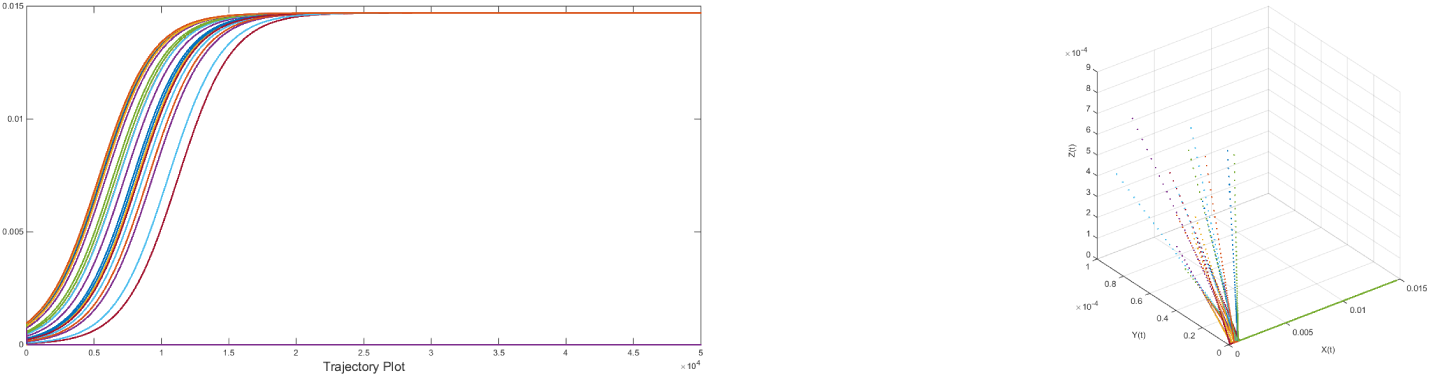
Left: Attracting trajectory of the fixed point(0.0147, 0, 0)for twenty different initial values(20 different set of colors are presented in the figure) taken from the neighbourhood of the fixed point, Right: corresponding three dimensional phase space.

Small immigration(*r*(*z*) = *rz*) relative to the density of the omnivore in the system as in Case *J* 13 leading to different dynamical behavior. There are five non-negative fixed points exist and one of which is not locally asymptotically unstable. One of the fixed points 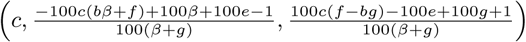 retains the coexistence of all the three species(prey, predator and omnivore) where all the parameters of the system involved. It is seen that only omnivore vanishes(prey and predator coexist) for the fixed point(*c,* 1 − *bc,* 0) and that existence of prey and predator depend on the carrying capacity of the prey and on the death rate of the predator in the absence of prey. On the other side, for the other fixed point 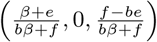 only prey and omnivore coexist. For the rest fixed point 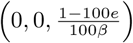 only omnivore exists. Only prey lives for the fixed points 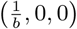.

#### Theorem 2.20.

*The fixed point(*0, 0, 0) *is asymptotically unstable for any non-negative parameters involved in the system.*

Here we put few examples of unstablity of the fixed point(0, 0, 0). Since the origin is unstable, the repelling trajectories yield to formation of chaotic and limit cycle attractors.

- (*b, c, e, f, g, β*) =(0.0546, 0.5013, 0.4317, 0.9976, 0.8116, 0.4857) then trajectories are slowly convergent to the fixed point(0.5013, 0.3037, 0.6689) as shown in the following Fig. 29 from top(1).
- (*b, c, e, f, g, β*) =(0.3119, 0.1790, 0.3390, 0.2101, 0.5102, 0.9064) then the origin repels and forms a limit cycle as seen in the phase space given in the following Fig. 29 from top(2).
- (*b, c, e, f, g, β*) =(0.2751, 0.2486, 0.4516, 0.2277, 0.8044, 0.9861) then the origin repels and forms a chaotic attractor(fractal dimension: 0.668, 0.783, 1.114) as seen in the phase space given in the following Fig. 29 from top(3).
- (*b, c, e, f, g, β*) =(0.0216, 0.9106, 0.8006, 0.7458, 0.8131, 0.3833) then the origin repels and forms a chaotic attractor(fractal dimension: 1.145, 1.783, 0.857) as seen in the phase space given in the following Fig. 29 from top(4).
- (*b, c, e, f, g, β*) =(0.0875, 0.6401, 0.1806, 0.0451, 0.7232, 0.3474) then the origin repels and forms a chaotic attractor(fractal dimension: 0.783, 0.897, 0.983) as seen in the phase space given in the following Fig. 29 from top(5).
- (*b, c, e, f, g, β*) =(0.1265, 0.1343, 0.0986, 0.1420, 0.1683, 0.1962) then trajectories forms a chaotic attractor(fractal dimension: 0.6874,0.65, 1.145) as seen in the phase space given in the following Fig. 29 from top(6).

## 3 Comparison among the Immigration and Classical Previte-Hoffman systems

When there is no immigration of any of the three species then the system is the classical Previte-Hoffman system which has been studied earlier. In this section, we shall compare of dynamical behaviour among the immigration and classical(non-immigration) Previte-Hoffman system through some specific examples.

**Figure 29:**
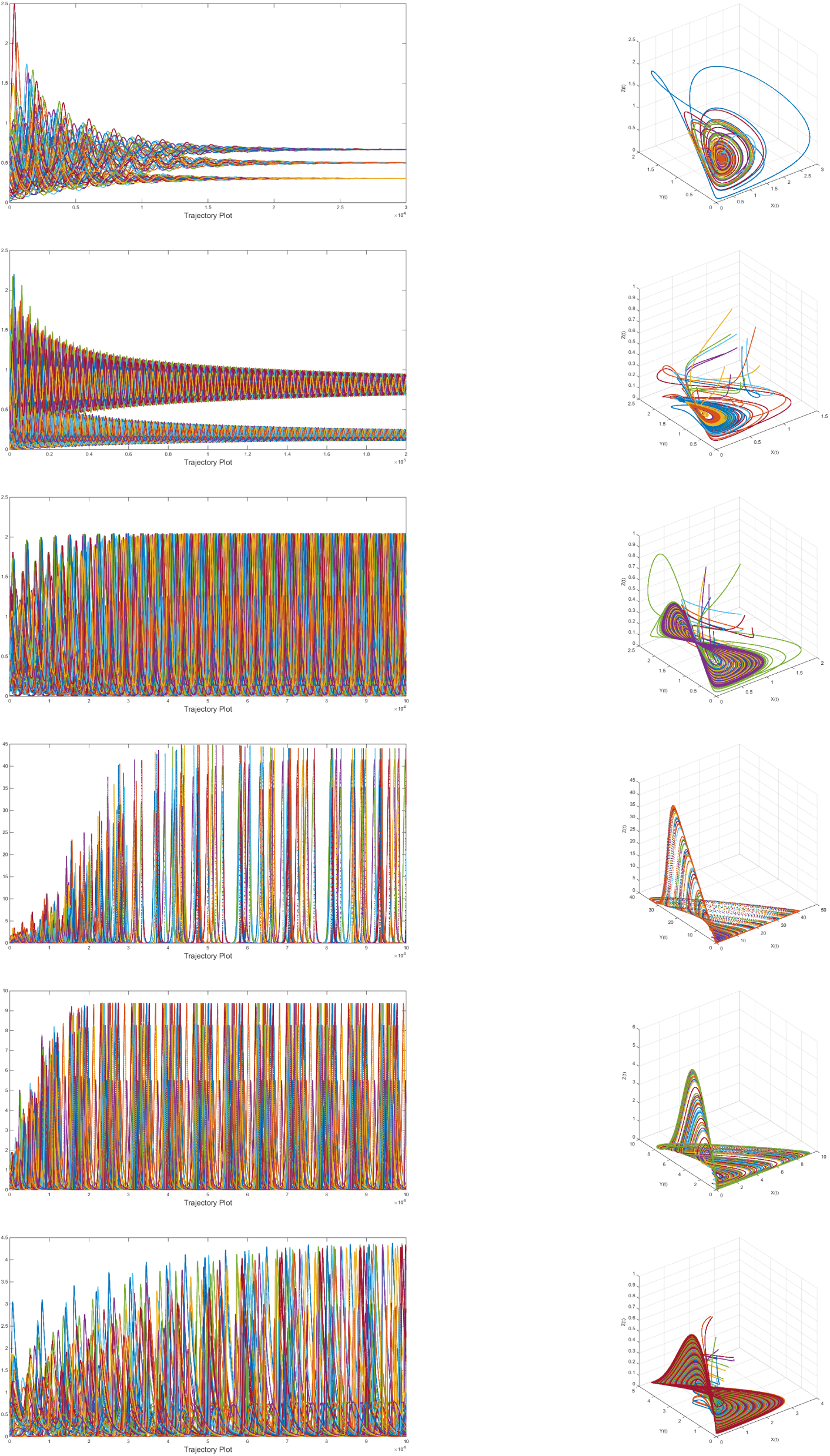
Left: Repelling trajectories from the fixed point(0, 0, 0) for twenty different initial values(20 different set of colors are presented in the figure) taken from the neighbourhood of the origin, Right: corresponding three dimensional phase space.

From the Table 4, it is observed that while transiting from the classical Previte-Hoffman model to Case I11 by introducing small(1%) immigration into the prey population, the fixed points got changed quantitatively but the qualitative behaviour(existence/coexistence of different population-prey, predator and omnivore) remain unchanged. Different color designate different quantitative behaviour. For all the three cases where 1% immigration are introduced in prey/predator/omnivore population then the origin is no more a fixed point while relative percentage of prey/predator/omnvore are introduced as adumbrated in all the three cases J11, J12 and J13 then origin remain a repelling fixed point as seen in previous section.

**Table 4:**
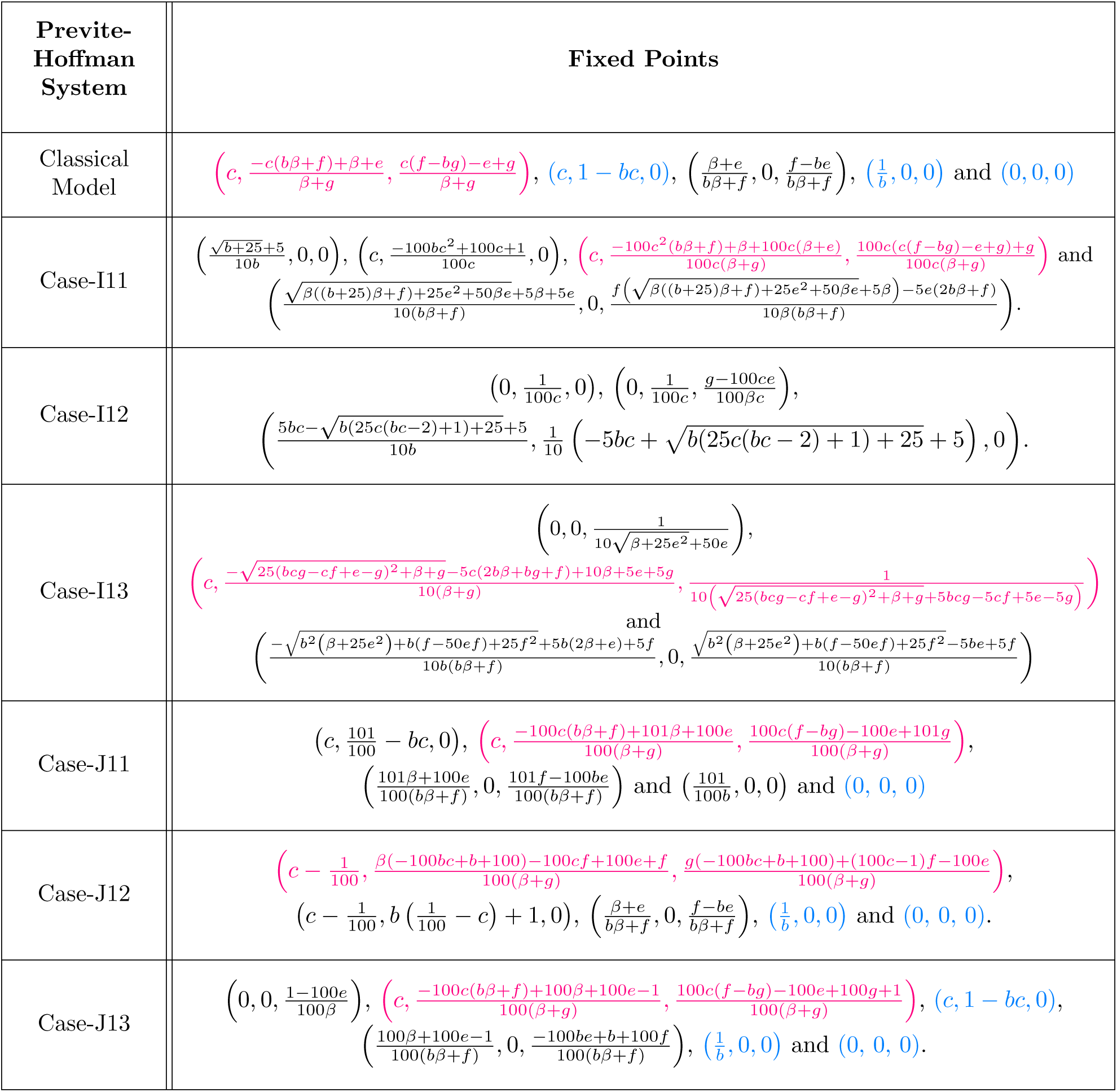
Fixed Points of the Immigration and Non-immigration Previte-Hoffman Systems.

Now we shall present a couple of examples for comparison among all these immigration and non-immigration Previte-Hoffman systems.

We take the parameters *b, c, e, f, g* and *β* as 0.1206, 0.5895, 0.2262, 0.3846, 0.5830 and 0.2518 respectively and the initial values for x, y and z are 0.8147, 0.9058 and 0.1270 respectively. The original Previte-Hoffman system exhibits chaos(fractal dimension: 0.986,01.125, 0.687). The trajectory plot is given in the Fig. 30.

**Figure 30:**
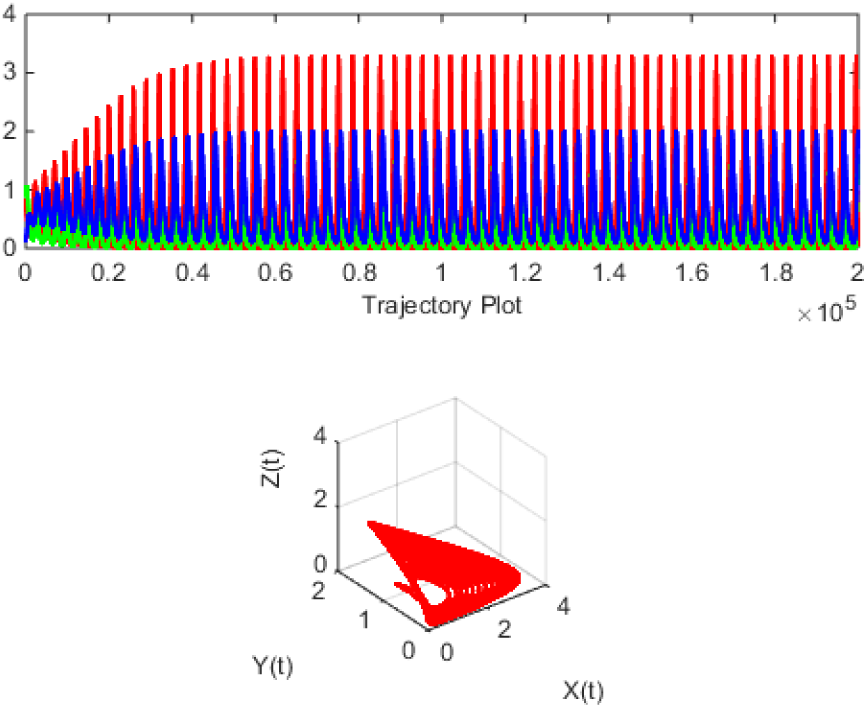
Up: Chaotic trajectory(x:red, y: green and z: blue) down: corresponding three dimensional phase space.

In all the six cases I11, I12, I13, J11, J12 and J13 the dynamics are given below and corresponding trajectory plot and phase are given in Fig. 31.

**Figure 31:**
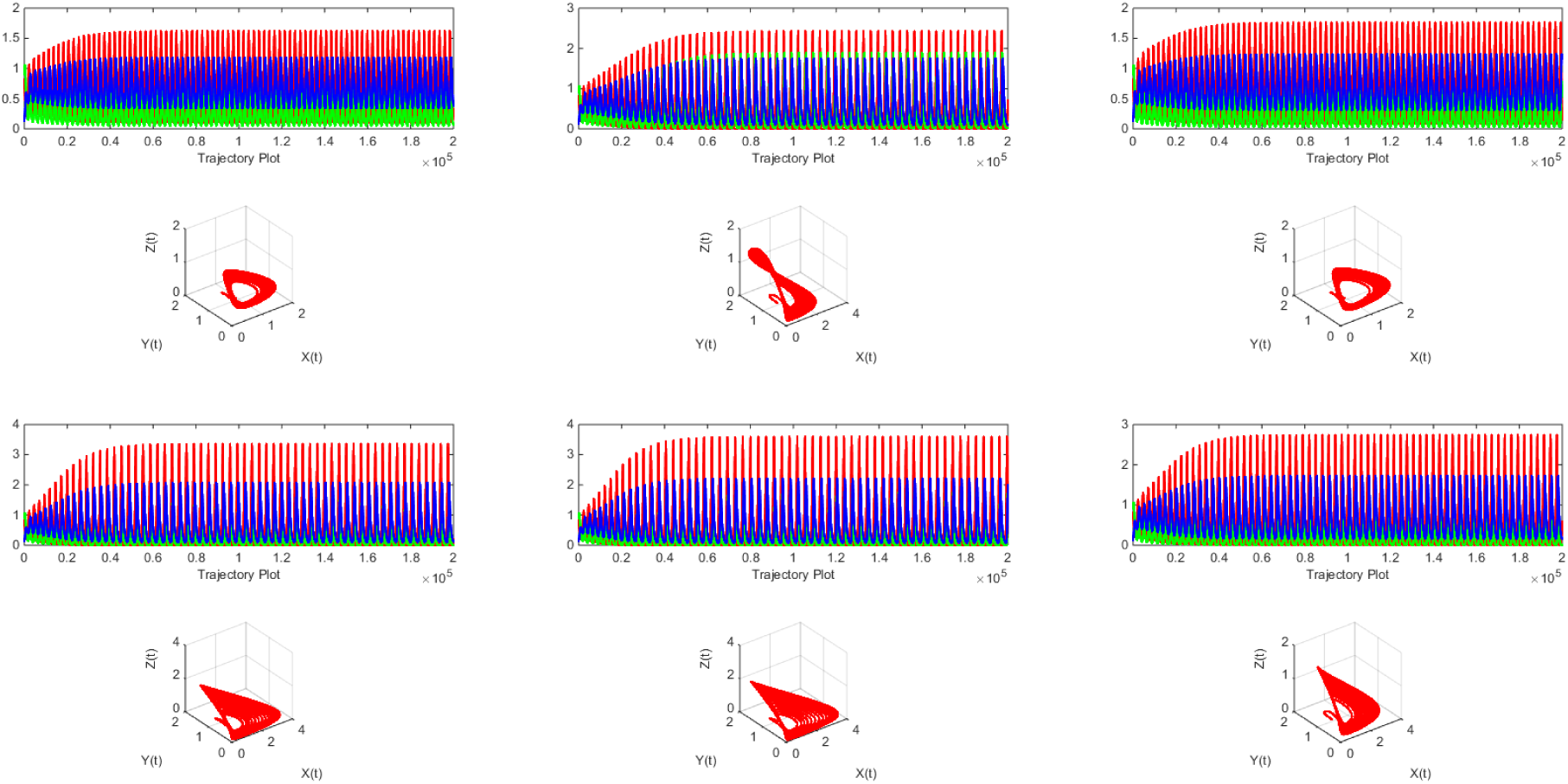
Trajectory plots and phase spaces for all starting from Case I11 to J13 from left to right(top to bottom)

**Case I11** For the fixed parameters and initial values, it is seen that the three dimensional trajectory possesses a limit cycle.

**Case I12** For the fixed parameters and initial values, it is seen that the three dimensional trajectory possesses a limit cycle.

**Case I13** For the fixed parameters and initial values, it is seen that the three dimensional trajectory possesses a limit cycle.

**Case J11** For the fixed parameters and initial values, it is seen that the three dimensional trajectory possesses a limit cycle.

**Case J12** For the fixed parameters and initial values, it is seen that the three dimensional trajectory possesses a limit cycle.

**Case J13** For the fixed parameters and initial values, it is seen that the three dimensional trajectory possesses a limit cycle.

It is observed that the limit cycles for all the cases including classical Previte-Hoffman system are getting small over time evolution as clearly depicted from the phase spaces.

We take another set of parameters *b, c, e, f, g* and *β* as 0.1206, 0.5895, 0.2262, 0.3846, 0.5830 and 0.2518 respectively and the initial values for x, y and z are 0.8147, 0.9058 and 0.1270 respectively. The original Previte-Hoffman system converge to the fixed point(0.8173, 0.3667, 0.0000). The trajectory plot is given in the Fig. 32.

**Figure 32:**
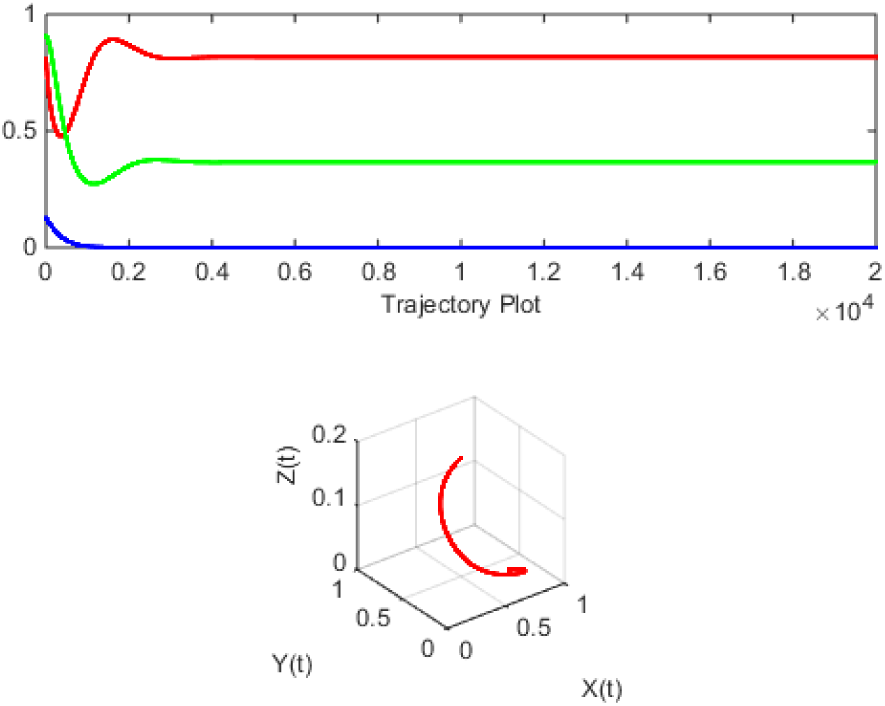
Up: Chaotic trajectory(x:red, y: green and z: blue) down: corresponding three dimensional phase space.

In all the six cases I11, I12, I13, J11, J12 and J13 the dynamics are given in the following Table 5 and corresponding trajectory plot and phase are given in Fig. 33.

**Figure 33:**
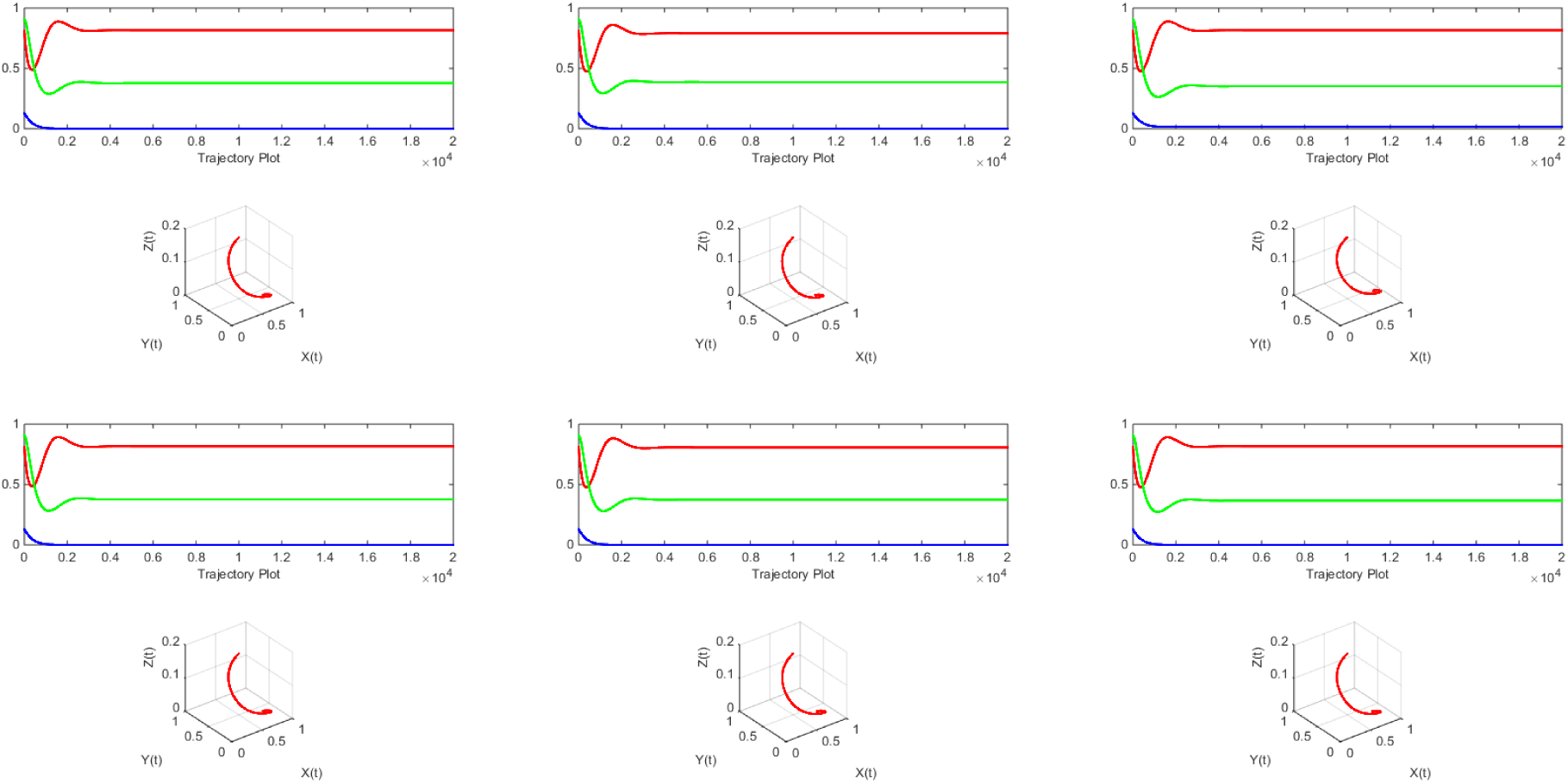
Trajectory plots and phase spaces for all starting from Case I11 to J13 from left to right(top to bottom)

**Case I11** For the fixed parameters and initial values, it is seen that the three dimensional trajectory converges to(0.8173, 0.3789, 0.0000).

**Case I12** For the fixed parameters and initial values, it is seen that the three dimensional trajectory converges to(0.7914, 0.3867, 0.0000).

**Case I13** For the fixed parameters and initial values, it is seen that the three dimensional trajectory converges to(0.8173, 0.3516, 0.0151).

**Case J11** For the fixed parameters and initial values, it is seen that the three dimensional trajectory converges to(0.8173, 0.3767, 0.0000).

**Case J12** For the fixed parameters and initial values, it is seen that the three dimensional trajectory converges to(0.8073, 0.3744, 0.0000).

**Case J13** For the fixed parameters and initial values, it is seen that the three dimensional trajectory converges to(0.8173, 0.3667, 0.0000).

It is noted that the qualitative behaviour of these six different cases of the system is remained unchanged with the original classical Previte-Hoffman system as seen above. It is observed that for the the cases J11, J12 and J13 where a relative percentage of immigration with respect to density of the respective population, the systems qualitatively behave like the original system as evident from the two examples cited.

## 4 Concluding Remarks & Future Endeavours

This article has presented different modified systems of the classical Previte-Hoffman model with notion of small immigration introduced into the different populations. It has been observed that the qualitative dynamics is remain unchanged. This is essentially establish the robustness of dynamics up to the small immigrations into its populations. This study suggests a detail quantitative and qualitative understanding of the asymptotic stability of the model in present of omnivore as well as immigration into different species involved. Such study needs experimental studies that estimate the relative contributions of the different resources in the omnivores diet and amount of immigration into the system. Further complicated biologically relevent immigration and migration can be introduced which we shall take up in our near future endeavour. Also the notion of immigration can be introduced into some other such ecological model in order to see the effect in the stability of those model.

## Acknowledgement

The author thanks to the *Pingla Thana Mahavidyalaya, Maligram, Paschim Medinipur, West Bengal* for generous support in executing the present work by providing the summer recess in 2018.

## References

1. Levin, S., 2003. Complex adaptive systems: exploring the known, the unknown and the unknowable. Bulletin of the American Mathematical Society, 40(1), pp.3–19.

2. Berryman, A.A., 1992. The Orgins and Evolution of Predator-Prey Theory. Ecology, 73(5), pp.1530–1535.

3. Yamamura, N., Higashi, M., Behera, N. and Wakano, J.Y., 2004. Evolution of mutualism through spatial effects. Journal of Theoretical Biology, 226(4), pp.421–428.

4. Hardin, G., 1960. The competitive exclusion principle. science, 131(3409), pp.1292–1297.

5. Hastings, A. and Powell, T., 1991. Chaos in a threespecies food chain. Ecology, 72(3), pp.896–903.

6. Gilpin, M.E., 1979. Spiral chaos in a predator-prey model. The American Naturalist, 113(2), pp.306–308.

7. Hastings, A. and Powell, T., 1991. Chaos in a threespecies food chain. Ecology, 72(3), pp.896–903.

8. Deng, B., 2004. Food chain chaos with canard explosion. Chaos: An Interdisciplinary Journal of Nonlinear Science, 14(4), pp.1083–1092.

9. Edelstein-Keshet, L., 1988. Mathematical models in biology (Vol. 46). Siam.

10. Morozov, A., Petrovskii, S. and Li, B.L., 2004. Bifurcations and chaos in a predator-prey system with the Allee effect. Proceedings of the Royal Society of London B: Biological Sciences, 271(1546), pp.1407–1414.

11. Beddington, J.R., Free, C.A. and Lawton, J.H., 1975. Dynamic complexity in predator-prey models framed in difference equations. Nature, 255(5503), p.58.

12. Tanabe, K. and Namba, T., 2005. Omnivory creates chaos in simple food web models. Ecology, 86(12), pp.3411–3414.

13. Walter, C., 1974. The global asymptotic stability of prey-predator systems with second-order dissipation. Bulletin of mathematical biology, 36(2), pp.215–217.

14. Wilmers, C.C. and Getz, W.M., 2004. Simulating the effects of wolf-elk population dynamics on resource flow to scavengers. Ecological Modelling, 177(1-2), pp.193–208

15. Coll, M. and Guershon, M., 2002. Omnivory in terrestrial arthropods: mixing plant and prey diets. Annual review of entomology, 47(1), pp.267–297.

16. Denno, R.F. and Fagan, W.F., 2003. Might nitrogen limitation promote omnivory among carnivorous arthropods?. Ecology, 84(10), pp.2522–2531.

17. Holt, R.D. and Polis, G.A., 1997. A theoretical framework for intraguild predation. The American Naturalist, 149(4), pp.745–764.

18. Gomes, A.A., 1998. A study of a three species food chain. Ecological Modelling, 2(110), pp.119–133.

19. Berryman, A.A., 1992. The Orgins and Evolution of PredatorPrey Theory. Ecology, 73(5), pp.1530–1535.

20. Barbosa, P. and Castellanos, I. eds., 2005. Ecology of predator-prey interactions. Oxford University Press.

21. Hassell, M.P., 1978. The dynamics of arthropod predator-prey systems. Princeton University Press.

22. Morin, P.J. and Lawler, S.P., 1995. Food web architecture and population dynamics: theory and empirical evidence. Annual Review of Ecology and Systematics, 26(1), pp.505–529.

23. Arim, M. and Marquet, P.A., 2004. Intraguild predation: a widespread interaction related to species biology. Ecology Letters, 7(7), pp.557–564.

24. Vance, R.R., 1978. Predation and Resource Partitioning in One Predator–Two Prey Model Communities. The American Naturalist, 112(987), pp.797–813.

25. Mehboob, M., Ahmad, S., Aqeel, M., Ahmed, F. and Ali, A., 2017. Turing bifurcation analysis for a predator-prey reaction-diffusion system. The European Physical Journal Plus, 132(9), p.399.

26. Owolabi, K.M., 2018. Modelling and simulation of a dynamical system with the Atangana-Baleanu fractional derivative. The European Physical Journal Plus, 133(1), p.15.

27. Huang, H.W. and Morowitz, H.J., 1972. A method for phenomenological analysis of ecological data. Journal of theoretical biology, 35(3), pp.489–503.

28. Tahara, T., Gavina, M.K.A., Kawano, T., Tubay, J.M., Rabajante, J.F., Ito, H., Morita, S., Ichinose, G., Okabe, T., Togashi, T. and Tainaka, K.I., 2018. Asymptotic stability of a modified Lotka-Volterra model with small immigrations. Scientific reports, 8(1), p.7029.

29. Previte, J.P. and Hoffman, K.A., 2013. Period doubling cascades in a predator-prey model with a scavenger. Siam Review, 55(3), pp.523–546.

30. Al-Khedhairi, A., Elsadany, A.A., Elsonbaty, A. and Abdelwahab, A.G., 2018. Dynamical study of a chaotic predator-prey model with an omnivore. The European Physical Journal Plus, 133(1), p.29.

31. Hofbauer, J. and Sigmund, K., 2003. Evolutionary game dynamics. Bulletin of the American Mathematical Society, 40(4), pp.479–519.

32. Hirsh, M.W., 1982. Systems of differential equations which are competitive or cooperative I: limit sets. Siam J. Appl. Math, 13, pp.167–179.

33. Hurwitz, A., 1964. On the conditions under which an equation has only roots with negative real parts. Selected papers on mathematical trends in control theory, 65, pp.273–284.

